# Control of Chloroplast Integrity by the Jasmonate Signaling Pathway is Linked to Growth-Defense Balance

**DOI:** 10.1101/2025.08.15.670541

**Authors:** Leah Y.D. Johnson, Ian T. Major, Qiang Guo, Yuki Yoshida, David M. Kramer, Gregg A. Howe

## Abstract

Chloroplasts play a central role in plant responses to environmental stress. Little is known, however, about how chloroplast homeostasis is maintained during stress responses that place high metabolic and bioenergetic demands on the cell. As a chloroplast-derived retrograde signal, jasmonate (JA) promotes broad-spectrum immunity by triggering the degradation of JAZ transcriptional repressors that act in the nucleus to control chloroplast metabolism. Here, we manipulated JAZ abundance to investigate how chloroplast integrity and function is maintained at high levels of defense. A *jaz* decuple mutant (*jazD*) lacking 10 of 13 JAZs exhibited strong growth-defense antagonism without loss of photosynthetic efficiency. Treatment of *jazD* with the JA-receptor agonist coronatine triggered rapid loss of chlorophyll and the turnover of chloroplast proteins and lipids, leading to the collapse of photosynthetic activity and cell death. These findings were supported by global transcript and metabolite profiling over a time course of coronatine treatment. Genetic screens identified MYC2 and the JAZ-destabilizing F-box protein, COI1, as positive regulators of coronatine-induced chloroplast dismantling in *jazD* plants. These results demonstrate how the progressive loss of JAZ repression drives a continuum of MYC2-dependent growth-defense tradeoffs, including disassembly of the photosynthetic apparatus as a terminal response. In highlighting the critical role for JAZ proteins in maintaining chloroplast integrity at high levels of defense, our results provide insight into the general mechanism by which jasmonate governs chloroplast metabolism to balance growth and stress responses.

**One sentence summary:** This study shows that the jasmonate-mediated leaf transition from growth- to defense-oriented metabolism culminates in the disassembly of the photosynthetic apparatus.

---

Photosynthesis converts inorganic compounds into complex macromolecules that fuel metabolism and growth. In plants, a major portion of photosynthetic products are sequestered in the chloroplast in the form of abundant proteins (e.g., Ribulose-1,5-bisphosphate carboxylase/oxygenase; rubisco), galactolipids, and metabolites such as chlorophyll. As leaves approach the end of their life span, they enter the senescence stage of development in which photosynthetic complexes are systematically dismantled into reusable metabolic precursors such as amino acids and fatty acids. The first visible symptom of senescence is the loss of chlorophyll (i.e., chlorosis), followed by turnover of photosynthetic proteins, unstacking of thylakoid membranes, and the proliferation of plastoglobules (Himelblau and Amasino, 2001; Kuai et al., 2018; Tamary et al., 2019; Domínguez and Cejudo, 2021). Metabolic products of senescence are mobilized to sink tissues and, in some cases, may be repurposed as alternative substrates for respiration (Yang and Ohlrogge, 2009; Araújo et al., 2011).

The onset of leaf senescence is triggered not only by age-dependent cues in mature leaves (i.e., natural senescence) but also by stress-related signals in developing young leaves. Among the factors that promote senescence in young leaves are energy starvation caused by light deprivation and disruption of reactive oxygen species (ROS) homeostasis (Liebsch and Keech, 2016; Domínguez and Cejudo, 2021). Stress-related hormones such as ethylene, abscisic acid (ABA), and jasmonate (JA) are well characterized for their role in promoting chlorophyll loss in young leaves (Lim et al., 2007). These hormones exert their effects in part by activating the expression of transcription factors (TFs), which in turn upregulate the expression of chlorophyll catabolic genes and other senescence-associated genes (SAGs) (Lim et al., 2007; Kuai et al., 2018).

In their classic study, Ueda and Kato employed detached oat (*Avena sativa*) leaf segments in a bioassay to identify methyl-JA (MeJA) as a senescence-promoting substance from *Artemisia absinthium* (Ueda and Kato, 1980). Subsequent work demonstrated that induction of chlorophyll loss by exogenous MeJA was associated with the degradation of rubisco and large-scale repression of photosynthesis-related genes (Parthier, 1990; Creelman and Mullet, 1997; Bilgin et al., 2010; Attaran et al., 2014; Gasperini and Howe, 2024). Studies performed with the model plant Arabidopsis (*Arabidopsis thaliana*) showed that JA-induced chlorophyll catabolism depends on the canonical signaling pathway comprised of the CORONATINE INSENSITIVE1 (COI1) receptor for jasmonoyl-L-isoleucine (JA-Ile) (He et al., 2002; Shan et al., 2011), members of the MYC family of TFs (Jiang et al., 2014; Qi et al., 2015; Zhu et al., 2015; Major et al., 2017; Song et al., 2017), and JAZ proteins that repress MYC activity (Yu et al., 2016; Monte et al., 2018; Guo et al., 2018b). The COI1-JAZ-MYC signaling cascade interacts with several additional positive and negative transcriptional regulators of senescence (Schommer et al., 2008; Kim et al., 2015; Qi et al., 2015; Zhu et al., 2015; Zhuo et al., 2020; Yang et al., 2023). The ability of JA-responsive TFs to recognize promoter elements of chlorophyll catabolic genes supports a model in which JA activates a transcriptional program for chlorophyll catabolism and subsequent turnover of photosynthetic complexes (He et al., 2002; Shan et al., 2011; Qi et al., 2015; Zhu et al., 2015; Song et al., 2017; Zhuo et al., 2020). Given that JA signaling promotes stress resilience through the upregulation of chloroplast pathways involved in the biosynthesis of antioxidant and defense-related compounds (Sasaki-Sekimoto et al., 2005; Pauwels et al., 2008; Guo et al., 2018a; Guo et al., 2022), a key unresolved question is how these anabolic pathways are appropriately balanced with JA-mediated catabolic processes that disrupt photosynthetic capacity and chloroplast integrity.

The biological relevance of JA as a signal for promoting leaf senescence remains unclear. The finding that natural leaf senescence in Arabidopsis is accompanied by an increase in endogenous JA levels and the expression of some JA biosynthetic genes initially suggested that JA is a signal for age-dependent senescence (He et al., 2002). Subsequent work, however, provided evidence that JA is not required for the initiation or progression of natural senescence in Arabidopsis (Seltmann et al., 2010a; Seltmann et al., 2010b). This latter view is consistent with the observation that Arabidopsis mutants defective in JA biosynthesis or signaling do not exhibit obvious phenotypes related to age-dependent senescence (He et al., 2002; Schommer et al., 2008; Seltmann et al., 2010b; Shan et al., 2011; Springer et al., 2016). Moreover, exogenous JA is largely ineffective in promoting leaf chlorosis in intact, wild-type Arabidopsis plants (Attaran et al., 2014; Major et al., 2017; Guo et al., 2018b). For this reason, studies of JA-induced senescence in Arabidopsis are performed using detached leaves floated in a JA-containing solution for several days in the dark (He et al., 2002; Shan et al., 2011; Zhu et al., 2015). Although convenient for molecular analyses, this experimental setup can impose starvation conditions that disrupt the natural senescence process in whole adult plants (Liebsch and Keech, 2016; Toyota et al., 2018; Wang et al., 2022; Weaver and Amasino, 2001).

In addition to promoting broad-spectrum resistance to biotic stress, JA is also recognized as a potent inhibitor of vegetative growth and seed production (Baldwin, 1998; Yan et al., 2007; Zhang and Turner, 2008; Wang et al., 2018; Guo et al., 2018b). These pleiotropic phenotypes implicate JA as a key plastid-derived signal for balancing growth- and defense-related processes (Guo et al., 2018a). In previous studies, we characterized a series of higher-order Arabidopsis *jaz* mutants with varying levels of growth-defense tradeoff (Campos et al., 2016; Guo et al., 2018b; Major et al., 2020; Guo et al., 2022; Johnson et al., 2023). A *jaz* decuple mutant (called *jazD*) deficient in 10 of the 13 *JAZ* family members is highly sensitive to exogenous JA and, intriguingly, displayed senescence-like symptoms in response to treatment with the COI1 receptor agonist coronatine (COR) (Guo et al., 2018b).

In the present study, we used intact wild-type and *jazD* plants grown under normal laboratory conditions to investigate the hypothesis that JA-triggered loss of chloroplast integrity is linked to leaf growth-defense balance. Our findings are consistent with a model in which the progressive loss of JAZ repression drives a continuum of MYC-dependent growth-defense tradeoffs, including disassembly of the photosynthetic apparatus as a terminal response. This work highlights a critical role for JAZ proteins in maintaining chloroplast integrity under elevated stress conditions and, in doing so, offers a biological explanation for the general phenomenon of JA-induced leaf senescence.

## Results

### Coronatine Treatment Promotes the Rapid Loss of Chloroplast Integrity in *jazD* Plants

We previously showed that treatment of *jazD* with coronatine (COR) results in leaf necrosis (Guo et al., 2018b). To better define the timing of this response and its relationship to chloroplast integrity, we monitored leaf chlorophyll content following the treatment of intact, soil-grown wild-type (WT) and *jazD* plants with a solution containing 5 μM COR (Fig. 1). COR treatment had no effect on chlorophyll levels in WT plants over the duration of the 96-h time course. Chlorophyll levels in COR-treated *jazD* leaves, however, began to decline within 24 h and were largely depleted by 72 h. This was accompanied by leaf necrosis and death by 96 h (Fig. 1, A and B). We used a chlorophyll fluorescence imaging system (Attaran et al., 2014; Cruz et al., 2016) as a non-invasive assay of COR-induced changes in photosynthetic capacity. COR treatment of *jazD* resulted in a steady decline in the efficiency of photosystem II (ɸ_II_), which closely tracked the reduction in chlorophyll content (Fig. 1, C and D; and Video S1). COR treatment had no obvious effect on ɸ_II_ in WT plants. These findings demonstrate that COR treatment results in rapid loss of chlorophyll and photosynthetic capacity in *jazD* but not WT plants.

**Figure 1.**
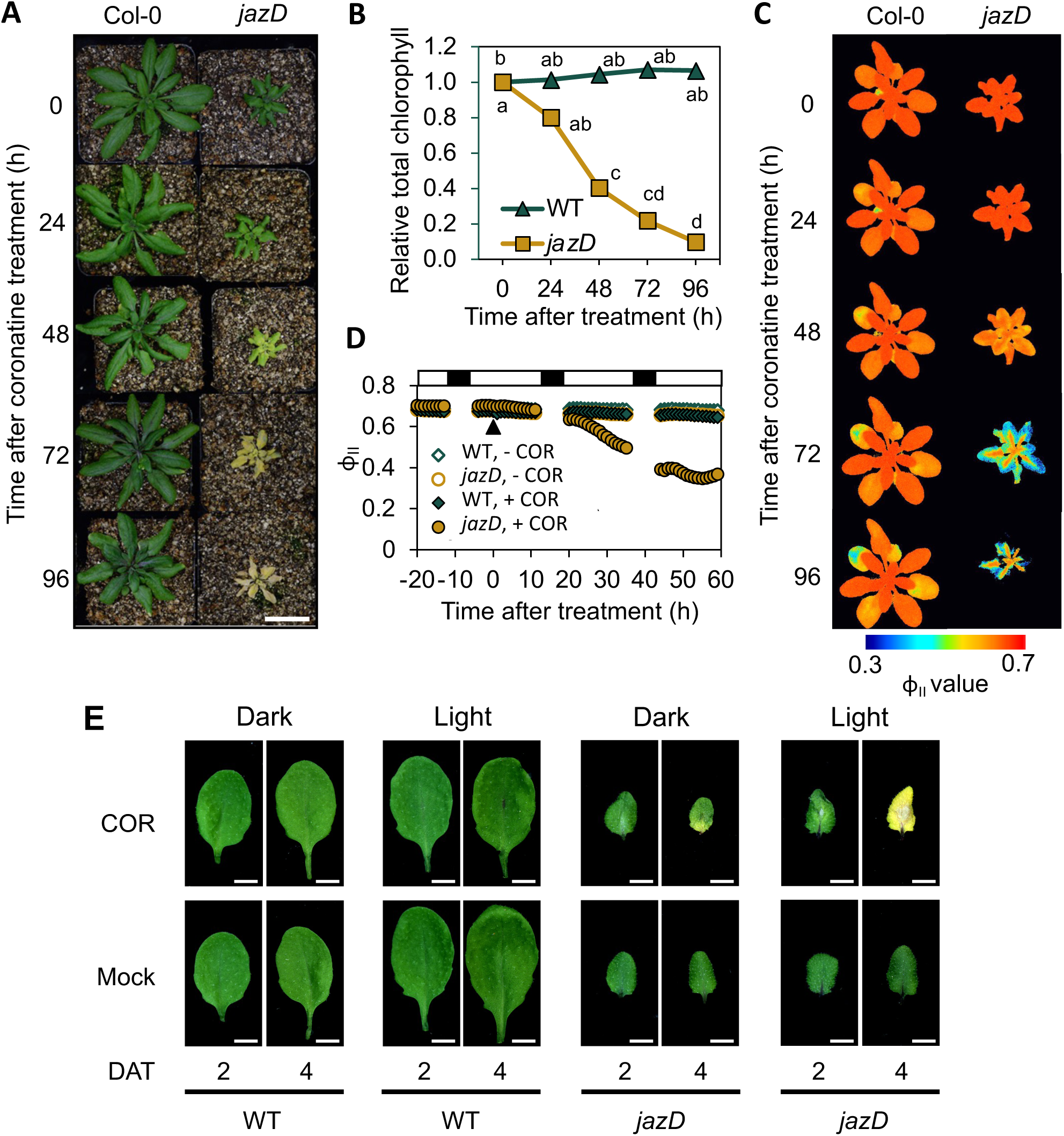
Coronatine promotes severe leaf chlorosis in *jazD* but not wild-type plants. A-B, WT and *jazD* plants grown in soil were sprayed with a solution containing 5 μM coronatine (COR). A, Plants with representative phenotypes at the indicated time point after treatment were photographed. Different plants were photographed at each time point. Scale bar, 2 cm. B, Quantification of total chlorophyll content in whole rosettes of WT and *jazD* plants after treatment with COR. Values were normalized to the amount of total chlorophyll at the “0” h time point. The data show the mean and SD of three plants per time point. Different letters represent significant differences at P<0.05 using Tukey’s honestly significant difference (HSD) post-hoc test. C-D, Coronatine treatment impairs photosynthetic efficiency in *jazD* but not WT plants. C, Soil-grown WT and *jazD* plants were acclimated in an imaging chamber for one day (negative time points in panel D) prior to treatment with COR. Vertically aligned images are representative false-colored images depicting the quantum efficiency of photosystem II (ɸ_II_) of the same plant, as measured by the Dynamic Environmental Photosynthetic Imaging (DEPI) system. The images were selected to match the time course shown in panel A. A time-lapse video of the experiment, including a WT and *jazD* plant treated with mock control, is available as Supplemental Video 1. D, Quantification of ɸ_II_ in WT and *jazD* plants treated with COR or mock control. The filled black triangle denotes the time of COR treatment. ɸ_II_ values were determined from DEPI images like those shown in panel C across the entire time course, with images acquired every hour and values averaged across the entire rosette. Bars at the top of the graph indicate light (white) and dark (black) cycles. E, COR-induced chlorosis in *jazD* leaves is strongly dependent on light. WT and *jazD* plants were grown under 16-h-light/8-h-dark photoperiod for 25 days. Five h after the beginning of the light cycle, the 5^th^ rosette leaf was spotted with 5 μL water (mock) or a solution containing 50 μM coronatine (COR). One set (Dark) of plants was immediately transferred to complete darkness for either 2 or 4 days, at which time the treated leaf was excised and photographed. A second set (Light) of COR- and mock-treated plants was kept under the normal photoperiod for 2 or 4 days. DAT, days after treatment. Scale bars, 0.3 cm.

Light plays a crucial role in regulating plant senescence by influencing both the timing and progression of chlorophyll degradation (Weaver and Amasino, 2001; Liebsch and Keech, 2016). We therefore tested the effect of normal growth light on COR-induced chlorosis of *jazD*. A single leaf from plants grown under standard photoperiod (16-h light/8-h dark) was treated with a 5 μl drop of COR or water (mock). Immediately following the treatment, plants were transferred to complete darkness for either two or four days. A control set of COR- and mock-treated plants was maintained under the normal light intensity and photoperiod. Based on visual leaf symptoms, we found that the severe chlorosis observed in COR-treated *jazD* was largely eliminated in plants that were transferred to the dark (Fig. 1E). Residual levels of chlorophyll loss in these dark-grown *jazD* plants, however, indicated that initiation of chlorosis by COR is not strictly dependent on light.

Chlorophyll catabolism during leaf senescence is typically accompanied by the degradation of abundant lipids and proteins that support photosynthesis (Lim et al., 2007; Woo et al., 2019; Tamary et al., 2019). To determine whether COR-induced senescence in *jazD* is associated with chloroplast lipid turnover, we quantified the levels of two prominent chloroplast lipids (monogalactosyldiacylglycerol, MGDG; and phosphatidylglycerol, PG) and one extra-plastidic lipid marker (phosphatidylethanolamine, PE) over a 3-d time course (Fig. 2A). The MGDG content in COR-treated *jazD* leaves decreased within 2 d of treatment, with levels declining to about 40% of the untreated control (“0” h time point) by day three. The abundance of PG in COR-treated *jazD* leaves followed a similar pattern, whereas the level of PE was not affected (Fig. 2A). COR treatment did not alter the relative abundance of MGDG, PG, or PE in WT plants.

**Figure 2.**
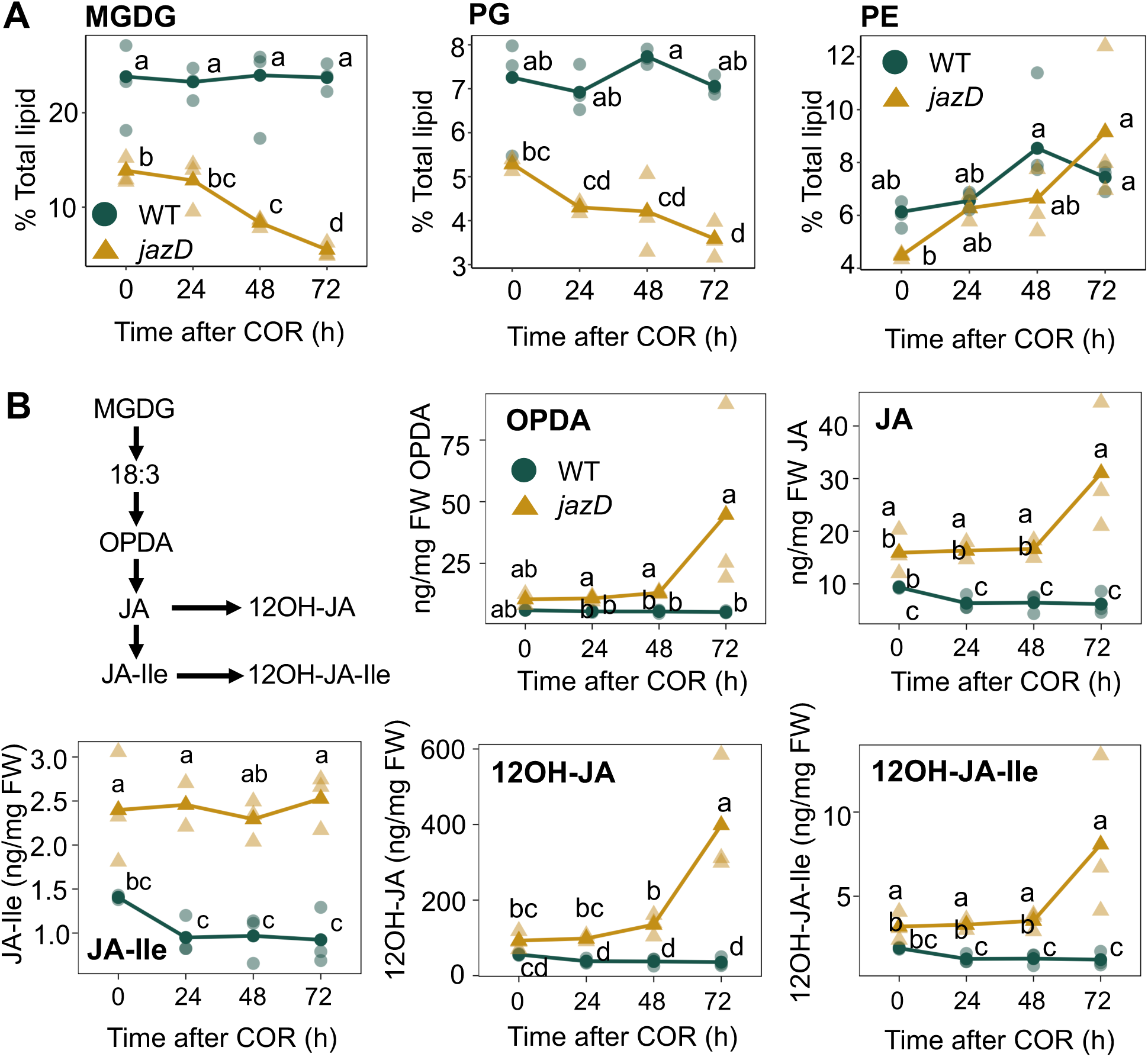
Coronatine treatment promotes the turnover of chloroplast lipids in *jazD*. A, Time course of polar lipid accumulation in response to coronatine treatment of wild-type (WT) and *jazD* plants. Intact plants were treated with a solution containing 5 μM coronatine and leaves were harvested for extraction and quantification of MGDG (monogalactosyldiacylglycerol; left panel), PG (phosphatidylglycerol; center panel), and PE (phosphatidylethanolamine; right panel) at the indicated time points after treatment. Low-case letters represent different statistical groupings from two-way ANOVA with Tukey’s HSD post-hoc test for each lipid (n = 3). B, Leaf jasmonate levels in response to coronatine treatment. WT and *jazD* plants were treated with a solution containing 5 μM coronatine and leaves were harvested for quantification of various jasmonate derivatives by LC-MS (n = 3). Different letters represent significant differences at P<0.05 using Tukey’s honestly significant difference (HSD) post-hoc test. OPDA, 12-oxophytodienoic acid; JA, jasmonic acid; JA-Ile, JA-isoleucine; 12OH-JA, 12-hydroxy-JA; 12OH-JA-Ile, 12-hydroxy-JA-Ile. Upper left panel depicts key steps in the biosynthesis and catabolism of JA-Ile from linolenic acid (18:3) moiety of MGDG lipids in the chloroplast.

We next asked whether COR-induced turnover of chloroplast lipids in *jazD* is associated with changes in the leaf jasmonate profile. COR treatment did not affect the abundance of 12-oxo-phytodienoic acid (OPDA), JA, JA-Ile, 12-hydroxy-JA (12OH-JA), or 12-hydroxy-JA-Ile (12OH- JA-Ile) in WT plants (Fig. 2B). In COR-treated *jazD* plants, however, the level of all JAs except JA-Ile trended higher at the 72-h time point. Among the largest effects in *jazD* was a ∼3-fold increase in the level of 12OH-JA in the third day of the time course (Fig. 2B). These data suggest that the COR-induced breakdown of chloroplast lipids in *jazD* generates precursors for the synthesis of JA and JA-Ile, which are subsequently converted to their corresponding 12- hydroxylated derivatives during the late stages of the time course.

Analysis of bulk leaf proteins by SDS-PAGE showed that COR treatment reduced the accumulation of major photosynthetic proteins in *jazD*, including the large and small subunits of rubisco and major light-harvesting complex proteins (Fig. 3A). Protein loss in COR-treated *jazD* leaves was evident within 48 h of treatment, with no obvious effects observed in WT leaves. We previously reported that *jazD* leaves overaccumulate an endoplasmic reticulum-localized β-glucosidase (BGLU18; AT1G52400) and cytosolic isoform of malic enzyme (ME2; AT5G11670), which co-migrate as a ∼65-kDa band (Guo et al., 2022). The presence of this band in COR-treated *jazD* remained relatively constant in comparison to rubisco (Fig. 3A), suggesting that COR treatment preferentially affects the accumulation of chloroplast proteins.

**Figure 3.**
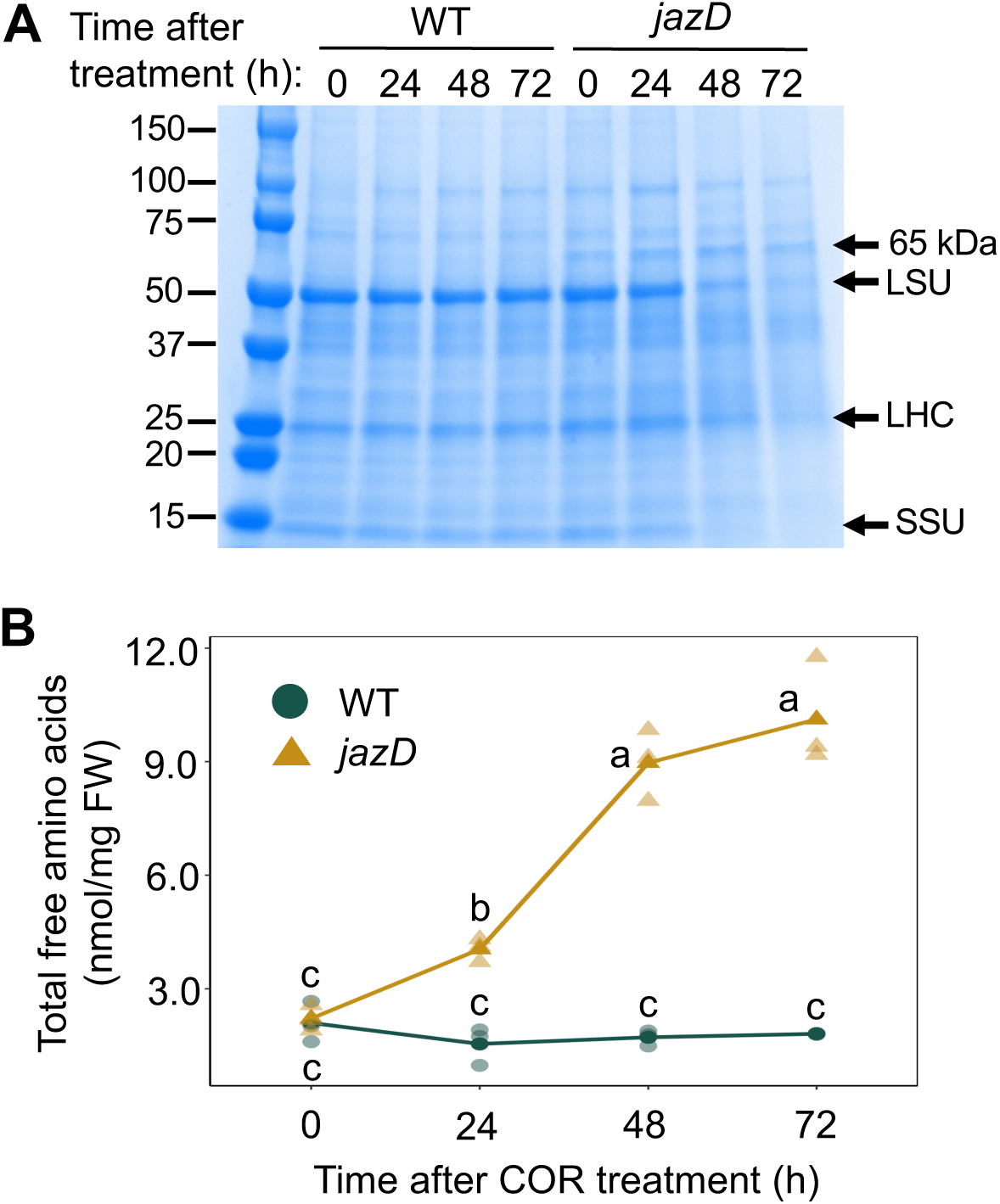
Time course of leaf protein turnover in response to coronatine treatment. Soil-grown wild-type (WT) and *jazD* plants were treated with a solution containing 5 μM coronatine and leaves were harvested for protein (A) or amino acid (B) extraction at the indicated time points. A, Proteins were separated on a Coomassie blue-stained SDS-PAGE (sodium dodecyl sulphate– polyacrylamide gel electrophoresis) gel. The large (LSU) and small (SSU) subunits of Rubisco and LIGHT HARVESTING COMPLEX II (LHC) are indicated, as is a 65-kDa, *jazD*-specific band previously shown to contain the extra-chloroplastic proteins β-GLUCOSIDASE 18 and Malic Enzyme 2 (Guo et al., 2022). Protein standards and their corresponding mass (kDa) are shown on the left. B, The 20 proteinaceous amino acids were individually quantified by LC-MS. The level of all 20 free amino acids was summed and plotted. Lines depict the mean value (solid symbol) of three biological replicates (opaque symbols). Different letters represent significant differences at P<0.05 using Tukey’s honestly significant difference (HSD) post-hoc test. The level of individual amino acids is shown in Supplemental Figure S1.

To further investigate how hyperactivation of jasmonate signaling impacts protein turnover, we quantified the level of proteinaceous amino acids in leaves of COR-treated WT and *jazD* plants. COR treatment had no effect on the amino acid content of WT leaves. The total level of free amino acids (i.e., summation of 20 proteinaceous amino acids) in *jazD*, however, increased 4.6-fold over the time course (Fig. 3B). This effect was accounted for by elevated levels of Ala, Arg, Asn, Cys, Gln, His, Ile, Leu, Lys, Met, Phe, Thr, Trp, Tyr, and Val (Fig. S1). Glu and Asp levels in *jazD* decreased during the time course, whereas Gly, Pro, and Ser levels remained relatively constant. These findings are consistent with a scenario in which COR-induced turnover of abundant chloroplast proteins in *jazD* contributes to the accumulation of free amino acids.

We next tested whether COR application alters the morphological features of chloroplasts in *jazD*. Light microscopy showed that COR treatment reduced the size of chloroplasts in leaf mesophyll cells of *jazD* within 72 h of application (Fig. 4A). Transmission electron microscopy further revealed that this response was associated with partial unstacking of thylakoid membranes (Fig. 4B). We also observed COR-induced proliferation of plastoglobuli, which is indicative of the developmental transition of chloroplasts to gerontoplasts in senescing leaves (Domínguez and Cejudo, 2021). COR treatment did not appear to affect the structural integrity of mitochondria in *jazD* (Fig. 4B). These collective data indicate that persistent activation of the JA signaling pathway in COR-treated *jazD* is associated with several biochemical and morphological hallmarks of leaf senescence.

**Figure 4.**
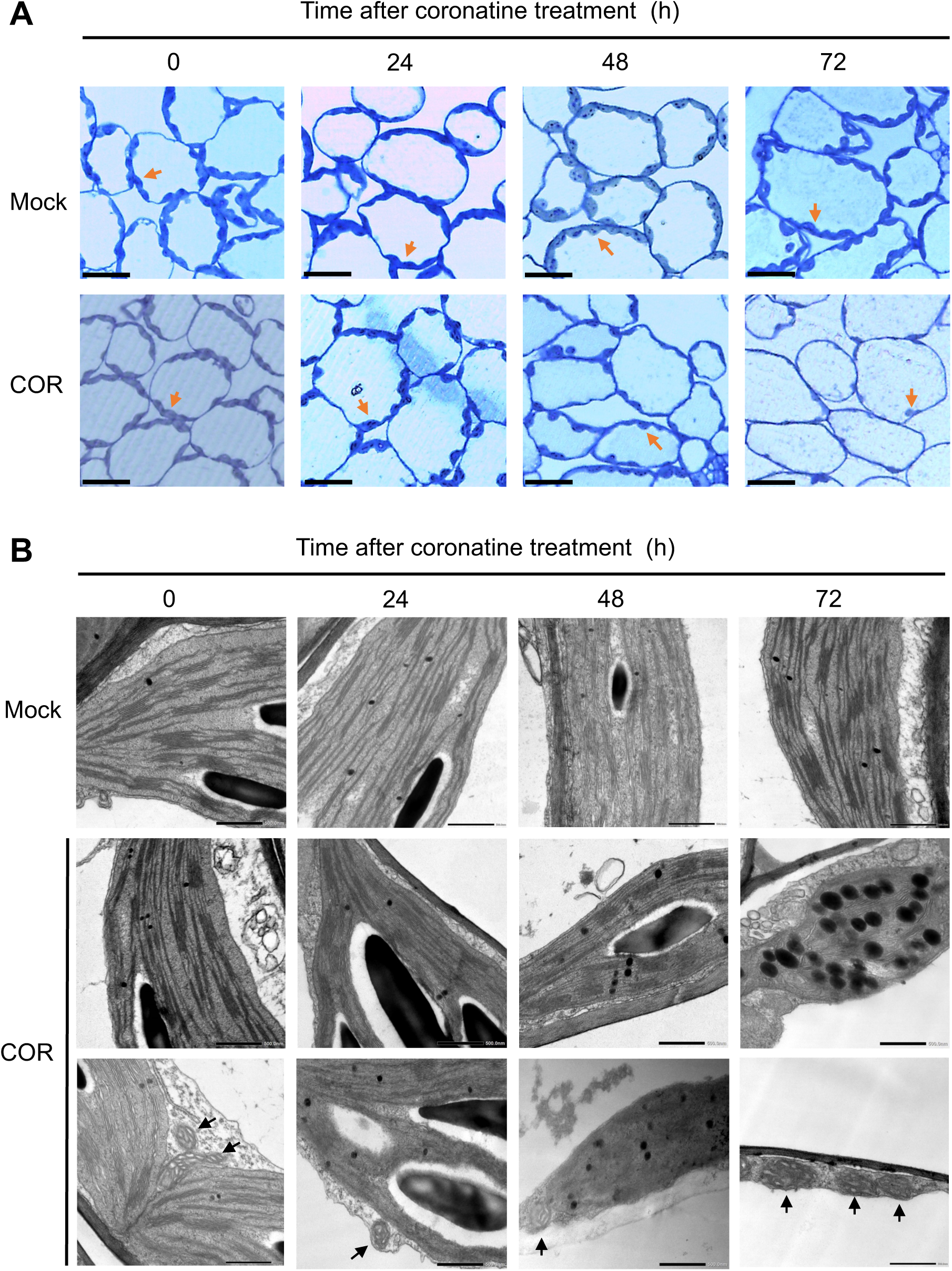
Coronatine treatment promotes the loss of chloroplast structural integrity in *jazD*. Twenty six-day-old *jazD* plants were treated with a solution containing 5 μM coronatine and leaves were harvested for imaging. A, Representative leaf cross-sections of *jazD*. Orange arrowheads indicate chloroplasts. Magnification is 400x, scale bar represents 20 μm. B, Transmission electron microscopy sections of representative chloroplasts and mitochondria from *jazD* plants treated with coronatine. Black arrowheads indicate mitochondria. Magnification, 15,000x; scale bar, 500 nm.

### Coronatine Treatment Elicits Defense- and Senescence-associated Gene Expression in *jazD*

To gain insight into how sustained JAZ deficiency reprograms transcriptional responses, we conducted an RNA-sequencing (RNA-seq) time-course experiment using RNA extracted from COR-treated *jazD* rosettes. Each time point (0, 1, 3, 12, and 24 h after COR application) included a mock-treated control, with three biological replicates per time point and treatment group (Fig. 5A). Transcripts whose abundance was ≥ 2-fold (log_2_) higher in the COR-treated sample relative to the time-matched mock control were classified as upregulated differentially expressed genes (DEGs). Similarly, mRNAs whose level was ≤ 2-fold (log_2_) lower in the COR-treated sample (relative to the matched mock control) were identified as downregulated DEGs. Further filtering for significant change in expression (P < 0.05) in COR- versus mock-treated samples generated a list of 7,676 genes that were differentially expressed at one or more time points (Supplemental Dataset 1). K-means clustering of the 7,676 DEGs identified six clusters representing distinct patterns of up- (clusters 2, 5, and 6) or down- (clusters 1, 3, and 6) regulated genes (Fig. 5B).

**Figure 5.**
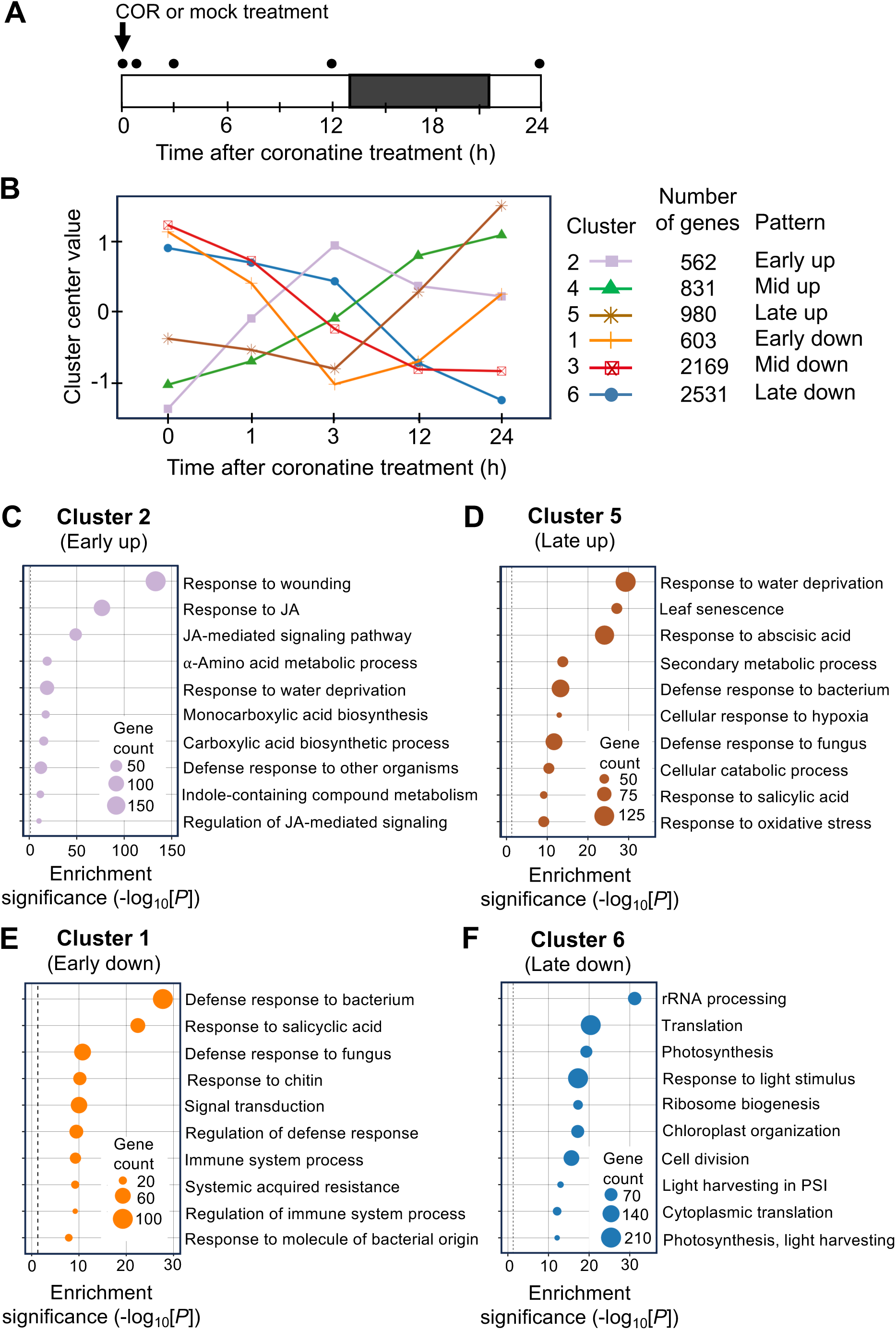
Global transcriptional response of *jazD* to coronatine treatment. A, Schematic of the RNA sequencing experiment. *jazD* plants were grown in soil under long-day photoperiod (open bar, light; black bar, dark) for 26 days and then sprayed with a solution containing 5 μM COR or a mock control. Whole rosettes were harvested at the indicated time points (circles) for RNA extraction using triplicate samples per time point for each treatment. B, RNA sequencing data was analyzed by *k*-means clustering to identify six groups of differentially expressed genes (DEGs) at each time point. The plot depicts *k*-means centroids for each cluster plotted over the time course. C-F) Gene ontology (GO) analysis (biological processes) was performed with genes comprising Clusters 2, 5, 1, or 6. Enriched functional categories were determined with DAVID using the hypergeometric test with Benjamini & Hochberg’s false discovery rate (FDR) correction.

We used the DAVID functional annotation tool (Sherman et al., 2022) to identify gene ontology (GO) terms that are enriched in each cluster (Supplemental Dataset 2). Cluster 2 (Early up) was highly enriched GO terms indicative of JA- and wound-mediated defense responses (Fig. 5C). Many of these are direct targets of MYC2; among 112 JA pathway genes shown to be bound by MYC2 (Zander et al., 2020), 45 (40%) of these were found in Cluster 2. This included genes encoding JA biosynthetic enzymes (LOX2/3/4, AOS, AOC, OPR3, OPCL1), JAZ repressors, and MYC2. Cluster 2 also included genes encoding JA oxidases (JOXs) and CYP94 P450s that negatively regulate the signaling pathway through catabolism of JA and JA-Ile.

GO categories within cluster 4 (Mid up) were indicative of wound- and JA-triggered processes involved in the biosynthesis of defensive secondary metabolites (Fig. S2 and Supplemental Dataset 2). Unlike cluster 2, cluster 4 also contained terms indicative of a senescence-associated shift from anabolic to catabolic processes. The upregulation of genes encoding proteases and chlorophyll catabolic enzymes (*PAO/ACD1, CLH1, and NYE2/SGR2*), for example, supported the biochemical phenotypes of COR-treated *jazD* plants. Consistent with the presence in cluster 4 of processes related to senescence, the abundance of *ORE1* transcripts encoding a major senescence-promoting transcription factor (Li et al., 2013; Qiu et al., 2015) increased steadily during the time course and reached peak levels at the 24 h time point (Supplemental Dataset 1). Transcripts in cluster 5 (Late up) exhibited relatively constant expression 3 h after COR treatment, followed by a sharp increase at later time points (Fig. 5D). This pattern was exemplified by additional chlorophyll catabolism genes (*NYC1* and *NYE1/SGR1*) as well as various senescence-associated genes. *SAG12*, which encodes a cysteine proteinase involved in nitrogen reallocation during senescence, is widely used as a marker of senescence (Noh and Amasino, 1999; James et al., 2018). Among all 7,676 DEGs identified in this RNA-seq experiment, *SAG12* exhibited the highest COR-induced expression at the 24 h time point (Supplemental Dataset 1).

Cluster 1 genes were sharply but transiently downregulated by COR treatment (Fig. 5). Enriched GO terms in cluster 1 were indicative of SA-mediated defense signaling (Fig. 5E and Supplemental Dataset 2) and associated signal transduction components, including transmembrane kinases and TFs. Specific examples of immunoregulatory genes that function in the SA branch of defense include *RPS2*, *PAD4*, *EDS1*, *SAG101*, *NPR4*, *WRKY53*, and *NDR1*. The downregulation of these genes during the time course is consistent with the well-established antagonism of SA responses by JA in Arabidopsis (Robert-Seilaniantz et al., 2011). Genes in cluster 3 showed a steady decrease in mRNA levels throughout the time course. Among the most significant GO categories in this group were “auxin-activated signaling”, “RNA modification”, and “plasma membrane” (Fig. S2 and Supplemental Dataset 2). The central role of auxin in promoting cell division and differentiation is consistent with the negative effect of JA signaling on growth-related processes (Noir et al., 2013). Cluster 3 also revealed the prolonged downregulation of nearly 100 genes encoding PENTATRICOPEPTIDE REPEAT (PPR) and TETRATRICOPEPTIDE REPEAT (TPR) proteins, many of which participate in RNA processing in plastids and mitochondria (Chi et al., 2008; Wang et al., 2021). Genes in cluster 6 were also persistently downregulated throughout the time course and, like cluster 3, was enriched in GO terms related to growth (Fig. 5F and Supplemental Dataset 2). A striking functional feature of cluster 6 was the preponderance of GO terms related to photosynthesis and ribosome assembly. The combination of genes comprising clusters 3 and 6 indicate that growth-related processes are strongly and persistently downregulated in COR-treated *jazD* plants.

### Coronatine-induced Senescence in *jazD* is Mediated by the Canonical COI1-JAZ-MYC2 Signaling Pathway

To investigate the molecular mechanism by which COR triggers leaf senescence in *jazD*, we performed a forward genetic screen for mutants that retain chlorophyll in response to COR treatment. We screened ∼16,000 ethyl methanesulfonate (EMS)-mutagenized *jazD* seedlings (M_2_ generation) for insensitivity to COR and identified ten putative *suppressor of coronatine hypersensitivity* (*sch*) mutants for further characterization (Fig. S3). Rescreening of these lines in the M_3_ generation confirmed the COR-resistant shoot phenotype and further showed that *sch* mutants were less sensitive than the *jazD* parental line to JA-induced root growth inhibition (Fig. S4).

Genetic complementation tests were performed to determine whether *sch* lines harbor mutations in either *COI1* or *MYC2*, which are required for many JA-induced responses in Arabidopsis. *sch* mutants were crossed to *jazD coi1* and *jazD myc2* tester lines (Fig. S5) and the resulting F_1_ seedlings were assessed for sensitivity to COR-induced chlorosis. Crosses between *jazD coi1* and *sch4*, *sch7*, *sch17*, and *sch19* yielded F_1_ progeny that retained resistance to COR, suggesting mutations in *COI1*. Sequencing of PCR-amplified genomic DNA from *sch4*, *sch7*, *sch17*, and *sch19* revealed the presence of G→A mutations that are predicted to generate single amino-acid substitutions in COI1 (Fig. 6A). Crosses between *jazD myc2* and *sch18*, *sch20*, *sch21*, *sch22*, *sch23*, and *sch24* produced F_1_ progeny that were all insensitive to COR-induced leaf chlorosis. Consistent with this finding, DNA sequencing of the *MYC2* gene from these six *sch* lines identified G→A nonsense mutations that are predicted to truncate the MYC2 protein (Fig. 6A). These data indicate that COR-induced leaf chlorosis of *jazD* is mediated by the canonical JA signaling pathway in which COI1 and MYC2 positively regulate the response.

**Figure 6.**
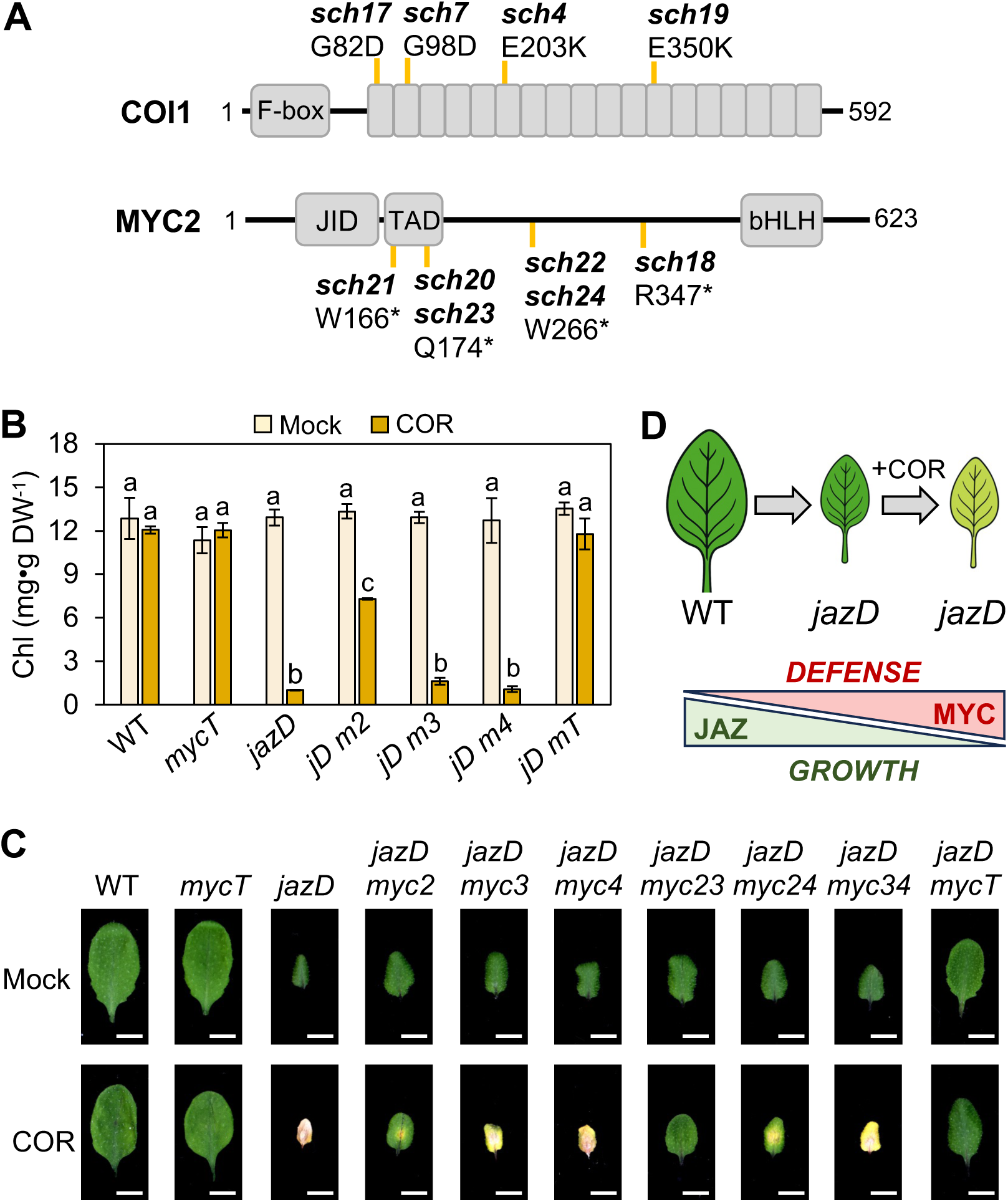
Coronatine-induced leaf chlorosis in *jazD* requires COI1 and MYC2. A, Identification of mutations in *COI1* (upper) and *MYC2* (lower) that impair coronatine (COR)-induced chlorosis in *sch* (*suppressor of coronatine-induced chlorosis*) mutants. The schematic diagrams depict the major domains of the Arabidopsis COI1 and MYC2 proteins, together with the position (vertical gold line) of *sch* point mutations mapped to the corresponding gene. All *sch* mutations are G-to-A transitions. Asterisks denote mutations that generate a premature stop codon. JID; JAZ-interacting domain; TAD, Transactivation domain; bHLH, basic helix-loop-helix domain. Small boxes in panel A correspond to the 18 leucine-rich repeats of COI1. B, Chlorophyll content in COR-treated leaves of select genotypes. Soil-grown plants (25-day-old) were sprayed with either water (mock) or a solution containing 5 μM COR. Leaves were harvested three days after treatment for chlorophyll quantification. Data show the mean ± SD of three samples per genotype. Letters denote significant differences according to Tukey’s HSD test (P < 0.05). *jD m2*, *jazD myc2*; *jD m3*, *jazD myc3*; *jD m4*, *jazD myc4*; *jD mT*, *jazD mycT*. C, The 5^th^ rosette leaf of 24-day-old plants of the indicated genotype was treated with a single 5 μL drop of water (mock) or a solution containing 50 μM COR. Leaves were excised and photographed four days after treatment. Scale bars, 0.5 cm. D, Schematic model (upper) depicting the progressive emergence of leaf growth inhibition and chlorosis phenotypes under three experimental conditions: WT (JAZ replete), *jazD* (JAZ deficiency), and COR-treated *jazD* (severe JAZ deficiency). These conditions lead to changes in the extent to which MYC transcription factors are repressed by JAZ proteins, which in turn modulates the balance between growth and defense.

*MYC2* and two closely related paralogs, *MYC3* and *MYC4*, have overlapping and redundant roles in a wide range of JA responses in Arabidopsis, including JA-mediated senescence of detached leaves (Fernández-Calvo et al., 2011; Qi et al., 2015; Zhu et al., 2015; Major et al., 2017; Johnson et al., 2023). To determine the contribution of each *MYC* paralog to COR-induced chlorosis, we quantified the effect of COR treatment on chlorophyll levels in *jazD* lines containing mutations in *MYC2/3/4*. As expected from the phenotype of *sch* mutants, *jazD myc2* plants exhibited strong (albeit partial) insensitivity to COR-induced chlorosis (Fig. 6B). Chlorophyll loss in *jazD myc3* and *jazD myc4* was indistinguishable from *jazD*, indicating that MYC3 and MYC4 are not required for the response in the presence of MYC2. The complete retention of chlorophyll in *jazD mycT*, which harbors a *myc2/3/4* triple mutation, suggested that *MYC3* or *MYC4* may contribute to COR-induced chlorosis in the absence of *MYC2*. Indeed, our qualitative leaf spotting assay provided evidence that *myc3* but not *myc4* enhances the COR-resistance of *jazD myc2* to the level observed in *jazD mycT* (Fig. 6C). We conclude that MYC2 performs the major role in promoting COR-induced loss of chloroplast integrity in *jazD*, with MYC3 contributing to chlorosis in the absence of MYC2.

## DISCUSSION

### Sustained JAZ Deficiency Activates Senescence-like Responses in Young Leaves of *jazD*

In this study, we addressed the question of how hyperactivation of the JA pathway through progressive loss of JAZ proteins affects chloroplast function and integrity. Our results show that treatment of young *jazD* plants with COR activates a series of molecular and physiological responses that resemble natural leaf senescence. The first visible symptom of COR application was leaf chlorosis, which was evident within 24 h of treatment. Chlorophyll loss was accompanied by the turnover of abundant chloroplast proteins and lipids, loss of photosynthetic efficiency, ultrastructural hallmarks of chloroplast disassembly, and tissue death. The strong (albeit partial) dependency of COR-induced leaf chlorosis on normal laboratory growth light is consistent with previous work showing that dark treatment of adult Arabidopsis plants delays natural senescence (Weaver and Amasino, 2001). Transmission electron microscopy provided evidence that mitochondria remained intact during COR-induced loss of chloroplast integrity in *jazD*. The apparent maintenance of mitochondrial integrity over the time course is noteworthy because mitochondria can provide an alternative source of cellular energy production during natural leaf senescence (Lim et al., 2007; Yang and Ohlrogge, 2009; Araújo et al., 2011; Woo et al., 2019). For example, it is possible that the COR-induced turnover of abundant chloroplast proteins and lipids in *jazD* provides substrates for mitochondrial respiration as a means of compensating for the loss of photosynthesis.

We found that the timing of leaf degreening in *jazD* was more rapid than typically observed for natural senescence. Whereas senescence in WT Arabidopsis unfolds over the course of weeks (depending on growth conditions) as plants naturally age (Lohman et al., 1994; Bresson et al., 2018), leaf chlorosis in intact *jazD* plants was visible within 24 h and complete within four days of COR treatment. Large-scale changes in senescence-associated gene expression were observed within 1 hour of COR application. This included induced expression of multiple SAGs (e.g., SAG12), as well as genes encoding chlorophyll catabolic enzymes. COR-triggered chlorosis was evident in all rosette leaves and thus appears to operate independently of leaf developmental age. The ability of COR treatment to synchronize a rapid senescence-like program throughout the *jazD* rosette may be useful for future studies aimed at identifying causal factors of JA-induced chlorosis and cell death.

### The Canonical JA Signaling Pathway Mediates Coronatine-induced Cell Death in *jazD*

We used an unbiased forward genetic approach to demonstrate that *COI1* and *MYC2* are required for COR-triggered lethality of *jazD*. The recovery of multiple, independent alleles of *coi1* and *myc2* suggests that the screen was saturated and thus that COI1 and MYC2 comprise major positive regulators of the response. Analysis of *myc2*, *myc3*, and *myc4* combinatorial mutants confirmed that MYC2 is the major contributor to COR-induced chlorosis. We also found that *MYC3* can promote chlorophyll loss in the absence of *MYC2*, in agreement with previous reports of functional redundancy between *MYC* paralogs (Qi et al., 2015; Zhu et al., 2015; Johnson et al., 2023). We conclude that the COR-induced phenotypes observed in intact *jazD* plants are mediated by the canonical COI1-JAZ-MYC2 signaling cascade. This finding supports previous studies of JA-induced senescence using excised leaf assays (He et al., 2002; Shan et al., 2011; Qi et al., 2015; Zhu et al., 2015; Major et al., 2017; Ding et al., 2022).

Our study highlights how changes in JAZ-MYC balance account for variation in the strength of growth-defense tradeoffs. In the absence of COR treatment, elevated MYC activity in *jazD* is sufficient to activate defense and restrict growth without negative effects on photosynthesis or other aspects of chloroplast function (Guo et al., 2018b; Guo et al., 2022; Johnson et al., 2023; this study). The dependency of COR-induced chlorosis on COI1 further indicates that the remaining pool of JAZ in *jazD* is degraded via the E3 ubiquitin ligase SCF^COI1^ and 26S proteasome. Turnover of these JAZs in *jazD* presumably further shifts the JAZ-MYC balance in favor of increased MYC activity. The ability of MYCs, together with downstream MYC-regulated TFs, to activate genes involved in chlorophyll degradation (Qi et al., 2015; Zhu et al., 2015; Kuai et al., 2018; Zander et al., 2020) provides a plausible explanation for the rapid onset of chlorosis following COR treatment. The resulting disruption of photosynthetic processes may also lead to the production of damaging reactive oxygen species that impair chloroplast integrity, as is the case during natural senescence (Bresson et al., 2018); this may explain our observation that COR-induced chlorosis is strongly dependent on light. These collective findings support a model in which the progressive loss of JAZ repression dynamically tunes growth-defense balance through increases in MYC activity that promote defense, restrict growth, and compromise chloroplast integrity at extreme levels of JAZ deficiency (Fig. 6D).

The absence of COR-induced chlorosis in WT plants (Fig. 1) likely reflects the existence of potent mechanisms to arrest JA signaling (Howe et al., 2018). For example, rapid JA-induced expression of *JAZ* genes in WT plants results in the de novo synthesis of JAZ proteins that are recalcitrant to JA-mediated degradation (Chung et al., 2010; Shyu et al., 2012; Moreno et al., 2013; Thireault et al., 2015; Zhang et al., 2017). The accumulation of transcripts encoding JAZ8, which is one of the few remaining JAZs in *jazD*, increased more than 30-fold within one hour of COR treatment. Nevertheless, it appears that the pool of JAZ repressors in *jazD* is insufficient to mitigate growth inhibition in the absence of COR treatment, and also insufficient to prevent more extreme phenotypes (e.g., chlorosis) following COR application (Fig. 6D). JA-mediated induction of metabolic pathways for catabolism of JA and JA-Ile provides another strategy to attenuate JA responses in WT plants (Koo et al., 2011; Heitz et al., 2019). Although our results indicate that JA and JA-Ile are converted to their hydroxylated derivatives in COR-treated *jazD* leaves, unique structural attributes of COR may render it resistant to these hormone-inactivating pathways.

MYC TFs are subject to multiple layers of regulation in addition to JAZ repression. *MYC2* mRNA abundance, for example, is positively regulated by the JA pathway (Lorenzo et al., 2004; Chung et al., 2008; Zander et al., 2020). We observed that *MYC2* transcript levels are upregulated (e.g., 8-fold increase at 3 h time point) in COR-treated *jazD* (Fig. S6). This positive feedback loop may contribute to the unrestrained JA responses in COR-elicited *jazD*. MYC TFs are also controlled by various post-translational mechanisms that influence the protein’s localization, stability, DNA binding capacity, and interaction with other TFs (Shin et al., 2012; Withers et al., 2012; Nakata et al., 2013; Schweizer et al., 2013; Zhai et al., 2013; Jung et al., 2015; An et al., 2017; Chico et al., 2020). Despite these multiple layers of control, our findings suggest that sustained loss of JAZ repression is sufficient to powerfully activate transcriptional responses mediated by MYC2.

### Coronatine Elicits Complex Transcriptional Responses in *jazD*

We found that COR treatment affects the expression of over 7,500 genes across the RNA-seq time course. This number represents approximately one-third of all leaf-expressed genes and supports previous analyses of leaf senescence in Arabidopsis (Breeze et al., 2011; Kim et al., 2016). The recovery of *coi1* mutations in the *jazD* suppressor screen indicates that these large-scale changes in gene expression are attributed specifically to hyperactivation of JA signaling rather than non-specific effects of exogenous COR. The finding that COR-induced loss of chloroplast integrity in *jazD* depends on MYC2 further demonstrates a critical role for this TF in shaping the underlying transcriptional response. Recent studies in tomato, rice, and Arabidopsis indicate that MYC2 orchestrates a hierarchical transcriptional cascade as part of the JA response (Du et al., 2017; Zander et al., 2020; Chen et al., 2024). We previously showed that unelicited *jazD* plants exhibit constitutive expression of multiple MYC-dependent responses, including reprogramming of primary and secondary metabolism, resistance to biotic stress, growth inhibition, and reduced seed yield (Guo et al., 2018b; Guo et al., 2022; Johnson et al., 2023). Here, we provide insight into how further de-repression of MYC TFs in *jazD* impacts global changes in gene expression.

COR treatment resulted in the rapid accumulation of nearly 1,400 transcripts represented by gene clusters 2 and 4. Comparison of our RNA-seq results to published ChIP-seq data (Zander et al., 2020) showed that clusters 2 (early up) and 4 (mid up) are enriched in targets of MYC2; 73% (411/562) of cluster 2 genes and 56% (463/831) of cluster 4 genes are directly bound by MYC2. Enrichment within these clusters of GO terms for DNA-binding TFs supports the view of MYC2 as a master regulatory factor that exerts its effects through a large network of secondary TFs (Du et al., 2017; Hickman et al., 2017; Zander et al., 2020). The 411 MYC2 targets in cluster 2 included 54 genes belonging to diverse TF families, including stress-related TFs in the MYB, ERF, bHLH, and NAC, and WRKY families. The likely role of these secondary TFs in executing COR-elicited responses in *jazD* is exemplified by several NAC TFs. *ANAC019/055/072*, for example, are direct MYC2 targets that control leaf senescence, including JA-induced chlorosis of excised leaves (Hickman et al., 2013; Zhu et al., 2015). These TFs bind to and activate the promoters of chlorophyll catabolic genes to initiate chlorosis (Zhu et al., 2015). Consistent with a model involving the temporal staging of primary (e.g., MYC2) and secondary TFs, the COR-induced expression of *ANAC019/055/072* appeared to precede increases in the level of mRNAs encoding chlorophyll catabolic enzymes (Fig. S6). We also observed rapid COR-induced expression of *ORE1*, which encodes a NAC TF with key roles in leaf aging and nutrient recycling (Fig. S6). *ORE1* expression is directly controlled by the EIN3 master regulator of ethylene responses (Chao et al., 1997; Qiu et al., 2015). Given that JAZs directly bind to and repress EIN3 action (Zhu et al., 2011), it is possible that COR-induced derepression of an EIN3-ORE1 module contributes senescence-like phenotypes in *jazD*.

In contrast to a detailed understanding of how the JA signaling pathway activates gene expression through relief-of-repression on JAZ targets such as MYC2 and EIN3, much less is known about the molecular mechanisms underlying JA-mediated downregulation of gene expression. Nearly 70% (5,303/7,676) of all differentially expressed genes in our RNA-seq experiment were categorized into one of three downregulated clusters (1, 3, or 6). Consistent with the notion that MYC2 functions mainly as a transcriptional activator (Zander et al., 2020), these downregulated clusters were not enriched in genes that bind MYC2. Nevertheless, there are examples of MYC2 target genes whose downregulation contributes to leaf senescence. In a study of JA-induced leaf senescence, MYC2 was shown to repress the expression of CATALASE 2 (CAT2), which in turn promotes senescence by elevating H_2_O_2_ levels (Zheng et al., 2020). Our RNA-seq data showing that *CAT2* mRNA levels in *jazD* are decreased in response to COR treatment supports this finding.

A noteworthy feature of downregulated genes in cluster 1 (early down) was the enrichment of GO terms for defense against bacteria and related immune system processes. This branch of plant immunity is dependent on SA and often exhibits antagonism with JA-mediated defense. Zheng and colleagues showed that a MYC2-ANAC019/055/072 regulatory module inhibits SA accumulation via the induced expression of the BSMT1 methyltransferase, which converts SA to volatile methyl-SA (Zheng et al., 2012). The massive increase in *BSMT1* mRNA levels (e.g., 500-fold 3 h post-COR treatment) in COR-treated *jazD* suggests that decreased SA levels may contribute to the repression of SA-mediated defense genes in *jazD*.

Enriched GO terms in clusters 3 and 6, which comprised most downregulated genes, were related to protein translation, ribosome biogenesis, cell division, and photosynthesis. The high cost of constructing and maintaining the cellular machinery for protein synthesis and photosynthesis is generally consistent with the active downregulation of these core growth-promoting processes during leaf senescence (Lim et al., 2007). Cellular Component GO terms within cluster 6 were dominated by chloroplast-related terms, including the chloroplast envelope membrane, thylakoid membrane, stroma, and thylakoid lumen. This observation is consistent with the gradual decline in the level of high abundance chloroplast proteins (e.g., rubisco), chloroplast lipids, and the upregulation of genes encoding chlorophyll catabolic enzymes. These findings support previous studies showing the activation of JA signaling globally downregulates the expression of photosynthesis-related genes (Bilgin et al., 2010; Attaran et al., 2014).

### Biological Significance of Jasmonate-induced Leaf Senescence

As a stress signal whose biosynthesis is initiated in the chloroplast, JA controls nuclear gene expression to modulate processes in many cellular compartments, including the chloroplast. This retrograde-like signaling circuit reprograms broad aspects of metabolism in response to specific environmental stress conditions. One of the most extensively studied metabolic effects of exogenous JA is chlorophyll catabolism and the associated turnover of abundant macromolecules in the chloroplast (Parthier, 1990; Hu et al., 2017). Interestingly, the efficacy of JA-induced chloroplast disassembly appears to depend on the plant species and experimental conditions. For instance, soil-grown (i.e., intact) rice and tobacco exhibit JA-induced chlorosis (Wang et al., 2022; Kim et al., 2023) whereas Arabidopsis does not (Fig. 1). For this reason, JA-induced senescence in Arabidopsis is typically studied by incubating detached leaves in a solution containing JA for several days in the dark (He et al., 2002; Shan et al., 2011; Zhu et al., 2015). This observation, together with the fact that JA biosynthesis and signaling mutants of Arabidopsis do not exhibit obvious defects in natural senescence, have raised questions about the physiological significance of JA-induced leaf senescence (Schommer et al., 2008; Seltmann et al., 2010a; Seltmann et al., 2010b).

Our results are consistent with the view that JA-induced leaf chlorosis (i.e., senescence) in young developing leaves is functionally linked to hyperactivation of JA-triggered immunity. It is well documented that plant immunity and senescence share various signaling components and physiological outputs, including nitrogen remobilization, ROS accumulation, and programmed cell death (Masclaux-Daubresse et al., 2010; Woo et al., 2019). Similar to JA, other plant stress hormones such as SA and ethylene work through master transcriptional regulators to simultaneously promote immune responses and senescence-like symptoms (Woo et al., 2019). These commonalities between senescence and immunity appear to reflect the role of stress hormones in controlling the strategic balance between survival and resource allocation.

JA-triggered immunity is energetically costly and requires an adequate supply of metabolic precursors to support the biosynthesis and storage of diverse defense-related compounds (Campos et al., 2014; Havko et al., 2016). Our findings support the view that JA-mediated turnover of chloroplast components reflects a nutrient remobilization process that is integral to the shift from growth- to defense-oriented metabolism. The major pool of cellular nitrogen is found in the chloroplast in the form of abundant photosynthetic proteins such as rubisco and light harvesting complexes (Evans and Clarke, 2019). The degradation of these proteins in COR-treated *jazD* leaves was associated with a rapid increase in the levels of most proteinogenic amino acids, suggesting that amino acid production from proteolysis outpaces the catabolism of these compounds. Interestingly, the level of some amino acids either remained unchanged (Ser, Pro, Gly) or declined (Glu and Asp) during the time course. The decline in Glu and Asp levels may reflect their use as amino donors, utilization as substrates in the TCA cycle, or entry into the GABA shunt (Miyashita and Good, 2008; Michaeli and Fromm, 2015). The steady accumulation of Ser and Gly may be linked to their roles in photorespiration and one carbon metabolism, whereas Pro homeostasis may be explained by its biosynthesis from Glu and oxidation in mitochondria.

A current limitation of our study is the lack of understanding of the extent to which JA-induced turnover of chloroplast components occurs in WT Arabidopsis. Despite the well documented role of MYC TFs in promoting chlorophyll catabolism and other aspects of chloroplast disassembly, we failed to detect a loss of chloroplast integrity in COR-treated WT plants grown under laboratory conditions. It is conceivable that increased MYC2 activity accelerates the basal turnover rate of abundant photosynthetic components as an integral part of the immune response of WT plants. Proteolysis of even a small fraction of abundant photosynthetic proteins (e.g., rubisco), for example, could provide a mechanism to rapidly to mobilize amino acids for the biosynthesis of defense compounds without compromising photosynthetic performance. It is also possible that the integration of JA signaling with other environmental inputs, such as changes in temperature and water availability, shape chloroplast metabolism in ways that were not detectable in our study (Havko et al., 2020). In this context, our findings highlight the dual role of chloroplasts as hubs for metabolic pathways that are integral to growth-defense balance and as potential reservoirs for nutrient recycling during plant responses to stress.

## MATERIALS AND METHODS

### Plant Materials and Growth Conditions

*Arabidopsis thaliana* accession Columbia-0 (Col-0) was used as the wild-type (WT) control for all experiments. Plants were grown in soil at 21°C under a long-day light regime of 16-h light (100 µmol m^-2^s^-1^) and 8-h dark. The construction of the *jaz* decuple (*jazD*), the *myc2 myc3 myc4* triple mutant (*mycT*), and *jazD myc* combinatorial mutants was previously described (Guo et al., 2018b; Guo et al., 2022; Johnson et al., 2023). The *jazD coi1* line used for genetic complementation assays was constructed as described in Supplemental Figure 5. Fluorescence measurements of photosynthetic parameters were performed with plants grown in the Dynamic Environmental Photosynthesis Imager (DEPI) as previously described (Attaran et al., 2014; Cruz et al., 2016). Plants were acclimated for 24 h in the DEPI chamber under non-fluctuating light conditions (100 µmol m^-2^s^-1^) prior to the acquisition of fluorescence data.

### Chlorophyll Measurements

Chlorophyll was extracted from whole rosettes (26 to 30-day-old plants) in 1 mL acetone:2.5 mM sodium phosphate (pH 7.8) (4:1 v/v) per 100-200 mg fresh weight (FW) tissue. Following centrifugation of the mixture for 10 min at 4°C, 800 µL of the resulting supernatant was transferred to a 1 cm cuvette and absorbances were measured at A_646_, A_663_, and A_750_. Chlorophyll concentrations were calculated as previously described (Porra et al., 1989) and the resulting values normalized to the volume of extraction buffer and tissue FW.

### Microscopy

Leaf sections from COR- and mock-treated *jazD* plants (26 to 29-day-old) were fixed in a solution containing 2.5% glutaraldehyde and 2.0% paraformaldehyde in 0.1 M cacodylate buffer. Samples were stained with 1% osmium tetroxide then fixed in Spurr resin (10.0 g 3,4-epoxycyclohexanemethyl 3,4-epoxycyclohexanecarboxylate; 6.0 g DER 736 Epoxy Resin; 26.0 g nonenyl succinic anhydride; 0.2 g 2-dimethylaminoethanol). Reagents described above were obtained from Electron Microscopy Sciences (EMS). Tissue sections used for transmission electron microscopy (TEM) were prepared using the Ultramicrotome Powertome XL (RMC Boeckeler) and placed on grids, followed by staining with 4% uranyl acetate and lead citrate as described (Reynolds, 1963). Light microscope sections were placed on glass slides and stained with Epoxy Tissue Stain (EMS). TEM images were acquired using a JEOL 1400 Flash Transmission Electron Microscope in the Center for Advanced Microscopy at Michigan State University. Light microscope images were acquired using a Leica DM750 microscope.

### Amino Acid Measurements

Amino acids were extracted with Milli-Q water containing ^13^C ^15^N-labeled amino acid standards (Sigma-Aldrich). Extracts were incubated at 90°C for 5 min and cooled on ice. The extracts were clarified by centrifugation and filtered through low-binding hydrophilic polytetrafluoroethylene (PTFE) filters (0.2 μM; Millipore). Filtered extracts were diluted with 20 mM perfluoroheptanoic acid (PFHA; Sigma-Aldrich) and analyzed using a Quattro Micro API LC–MS/MS (Waters) equipped with an Acquity 623 UPLC HSS T3 1.8 μm column (2.1 × 100 mm; 1.8 μm particle size; Waters) as described previously (Major et al., 2020). Amino acid concentrations were determined from an external standard curve and normalized to tissue fresh weight.

#### Hormone Measurements

WT and *jazD* plants were grown for 26 days and then treated with a solution containing 5 µM COR. Samples were extracted at 4°C overnight (∼16 h) using 0.45 ml ice-cold methanol:water (80:20 v/v) containing 0.1% formic acid, 0.1 g l−1 butylated hydroxytoluene (BHT) spiked with abscisic acid (ABA)-d6 (100 nM) as an internal standard. Filtered plant extracts were injected onto an Acquity UPLC BEH C18 column (2.1 x 100 mm; 1.7 µM particle size) and analyzed as described previously (Liu et al., 2016). Selected ion monitoring (SIM) was conducted in the negative electrospray channel for jasmonic acid (JA; *m*/*z* 209.1>59), JA-isoleucine (JA-Ile; *m*/*z* 322.2>130.1), 12-hydroxy-JA (12OH-JA; *m/z* 225.2>59), 12-hydroxy-JA-Ile (12OH-JA-Ile; *m/z* 338.2>130.1), 12-oxo-phytodienoic acid (OPDA; *m/z* 291.2>165.1) and the internal ABA-d6 standard (*m*/*z* 269.1>159.1). Hormone concentrations were determined from external standard curves and normalized to tissue fresh weight.

#### mRNA Sequencing and Analysis

Gene expression profiling was performed on the BGISEQ platform at BGI Genomics. Rosette leaves of 26-day-old soil-grown *jazD* plants were sprayed with a solution containing 5 μM COR or mock control 3 h after the start of the light cycle. The entire rosette was harvested at 0, 1, 3, 12, and 24 h after treatment and immediately frozen in liquid N_2_. For the 0 h time point, the rosette was harvested immediately after spraying (i.e., within 2-3 min of treatment) with mock or coronatine solutions. Whole rosettes from four individual plants were pooled for each sample. The tissue was homogenized using the TissueLyser II (Qiagen) and stored at -80C until further use. RNA was extracted using the NucleoSpin RNA Plant kit (Machery Nagel). RNA quality was assessed using the Agilent 4200 Tapestation and RNA concentrations were measured using the Qubit fluorometer (Invitrogen) in the MSU Genomics Core Facility. Three independent RNA samples (biological replicates) were used for each time point and treatment, with each replicate derived from pooling rosette leaves from four plants as described above.

Sequencing and data preprocessing was performed by BGI Genomics to provide adaptor-trimmed, high-quality reads. Raw sequencing reads were mapped to TAIR10 gene models by RSEM (version 1.3.3) (Li and Dewey, 2011). mRNA abundances for all Arabidopsis genes were expressed as transcripts per million (TPM). DESeq2 (version 3.12) was used to normalize expected counts from RSEM and to determine differential gene expression by comparing normalized counts in mock-treated samples to the coronatine-treated samples at each time point (Love et al., 2014). DAVID (version 2021) was used to perform gene ontology (GO) analysis of enriched functional categories (Sherman et al., 2022). Over- and underrepresented GO categories among differentially expressed genes were assessed by hypergeometric test with Benjamini & Hochberg’s False Discovery Rate (FDR) correction at P < 0.05.

#### Lipid Analysis

Monogalactosyldiacylglycerol (MGDG), phosphtidylglcerol (PG), and phosphatidylethanolamine (PE) were extracted, purified by thin layer chromatography, and analyzed by gas chromatography (GC) as previously described (Wang and Benning, 2011). Peak areas for individual fatty acyl methyl esters (16:0, 16:1 *cis* and *trans*, 16:2, 16:3, 18:0, 18:1, 18:2, and 18:3) of each lipid were determined. The percent of total lipid was calculated by summing the total abundance of each of the fatty acyl methyl esters for each lipid. This value was then divided by the total fatty acyl methyl ester abundance of the total lipid extract for each sample (Wang and Benning, 2011).

#### Protein Analysis

WT and *jazD* plants were grown in soil for 26 days and then treated with a solution containing 5 µM coronatine. Leaves harvested 0, 24, 48, and 72 h after COR treatment were flash-frozen in liquid nitrogen and stored at -80 °C. Proteins were extracted from equal amounts of fresh tissue in a buffer containing 80 mM Tris-HCl pH 6.8, 10% glycerol, 2% sodium dodecyl sulfate (SDS), and one tablet of Complete Mini ethylene diamine tetraacetic acid (EDTA)-free proteinase inhibitors (Roche) and 500 µL 2-mercaptoethanol per 10 mL buffer. Extracted proteins were separated by SDS–polyacrylamide gel electrophoresis (SDS-PAGE) (4-20% Mini-PROTEAN TGX Precast Protein Gel; BioRad) and stained with Coomassie Brilliant Blue R-250. Protein molecular weight standards were obtained from the All Blue Prestained Protein Standard (BioRad).

#### Selection of Coronatine-resistant Mutants

To generate the screening population, *jazD* seeds were treated for 16 h with a solution containing 0.1% EMS (ethyl methanesulfonate). The resulting M_1_ plants were grown for production of M_2_ seed. M_2_ seeds were pooled into 24 families and independent families were screened. For selection of COR-resistant mutants, M_2_ seeds were surface sterilized by treatment for five minutes in 70% (v/v) ethanol, five minutes in 30% (v/v) bleach, and rinsed in sterile deionized water six to eight times. Seeds were then stratified at 4°C for three to four days. Sterile seeds were plated onto media containing half-strength Linsmaier and Skoog (LS; Caisson Labs), 0.8% (w/v) sucrose, and 0.8% (w/v) Phytoblend agar (Caisson Labs). Plates were incubated under 16 h light (80 µmol m^-2^s^-1^) and 8 h dark conditions at 21°C. After eight days of growth, seedlings were sprayed with a solution containing 1 µM COR (Sigma Aldrich). Four days after COR treatment, seedlings were visually screened for reduced chlorosis and growth inhibition compared to the *jazD* control. Approximately 17,500 seeds were screened from eight independent families. To test coronatine resistance in mature plants, leaves of 26-d-old plants were spotted with 5 µl of a solution containing 50 µM COR. Leaves were excised and photographed four d after treatment. COR stocks were dissolved in 100% ethanol and diluted in water.

#### Sanger Sequencing for Identification of Mutations in *COI1* and *MYC2*

PCR amplification of genomic DNA containing *COI1* and *MYC2* from *sch* mutants was performed with primer sets spanning the 5’ and 3’ UTRs. Primer sequences are provided in Table S1. PCR reaction conditions were as follows: 95°C for 5 min, 35 cycles of denaturation (95°C, 30 s), annealing (59.6°C, 30 s for *MYC2*; 48.6°C, 30 s for *COI1*), elongation (72°C, 1.5 min), and one final elongation step at 72°C for 5 min. Reactions were held at 12°C until removed from the PCR instrument and analyzed on a 1% agarose gel with DNA standards (1 kb plus DNA ladder, Invitrogen). PCR products were purified from agarose gel slices using the Qiaquick Gel Extraction Kit (Qiagen). Purified product was combined with forward and reverse primers (Table S1) and sequenced at the MSU RTSF Genomics Core Facility. Sequencing reads were aligned to the genomic sequence to identify polymorphisms.

## Data availability

RNA sequencing raw and processed data for this study were deposited in the Gene Expression Omnibus (GEO) database (https://www.ncbi.nlm.nih.gov/geo) as accession no. GSE266991. Data available includes the raw paired-end reads for each sample, a CSV file containing the transcripts per million (TPM) for each sample, and an XLSX file containing the log2 fold-change expression data for each sample.

## Gene accession numbers

Genes described here have the following Arabidopsis Genome Initiative (AGI) gene accession numbers: *COI1* (AT2G39940), *JAZ1* (AT1G19180), *JAZ2* (AT1G74950), *JAZ3* (AT3G17860), *JAZ4* (AT1G48500), *JAZ5* (AT1G17380), *JAZ6* (AT1G72450), *JAZ7* (AT2G34600), *JAZ9* (AT1G70700), *JAZ10* (AT5G13220), *JAZ13* (AT3G22275), *MYC2* (AT1G32640), *MYC3* (AT5G46760), and *MYC4* (AT4G17880).

## Acknowledgements

We thank Nate Havko for technical advice throughout this study. We also acknowledge Jeff Cruz for expert assistance with fluorescence imaging, Ron Cook and Anastasiya Lavell for technical assistance with lipid measurements, and Tony Schilmiller in the MSU Mass Spectrometry Facility for helpful assistance with LC/MS/MS analysis of metabolites.

## Author contributions

L.Y.D.J. and G.A.H. conceived the project. L.Y.D.J., I.T.M., Q.G., and Y.Y., performed the experiments. D.M.K. assisted with the design of the chlorophyll fluorescence imaging experiments. L.Y.D.J. and G.A.H. wrote the paper with input from all authors. G.A.H. agrees to serve as the author responsible for contact and ensures communication.

## Funding information

This study was supported primarily by the Chemical Sciences, Geosciences, and Biosciences Division, Basic Energy Sciences, Office of Science at the U.S. Department of Energy (grant no. DE–FG02–91ER20021). The publication was made possible by a predoctoral training award to L.Y.D.J. from grant number T32-GM110523 from National Institute of General Medical Sciences of the National Institutes of Health. We also acknowledge support from Michigan State University AgBioResearch and a Dissertation Completion Fellowship to L.Y.D.J. from the College of Natural Science at Michigan State University.

## Notes

### Competing Interest Statement

The authors have declared no competing interest.

