## Supplemental material for "Control of Chloroplast Integrity by the Jasmonate Signaling Pathway is Linked to Growth-Defense Balance"

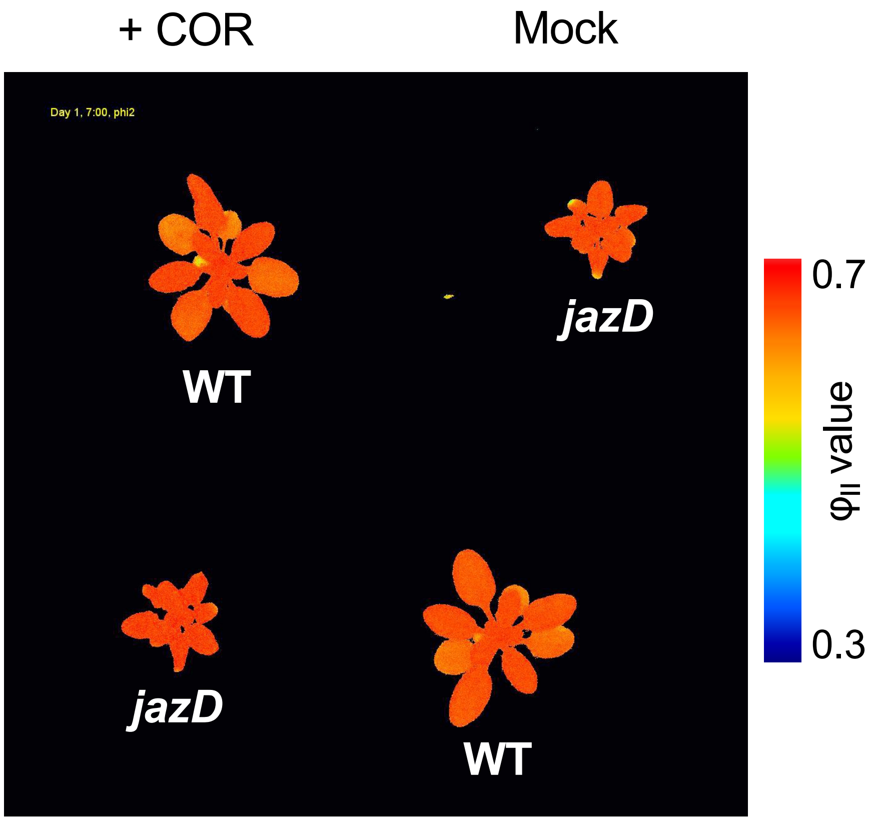


**Supplemental Video 1.** Time course of changes in photosystem II efficiency in response to treatment with coronatine. Three-week-old wild-type (WT) and *jazD* plants were grown in soil under a long-photoperiod conditions (16-hour day/8-hour night). Plants were transferred to a fluorescence imaging chamber and acclimated for 1 day (Day 1). Following this acclimation period, plants were sprayed (at 10 AM on Day 2) with either water (Mock; two plants at right) or a solution containing 5 uM coronatine (+COR; two plants at left). Photosystem II efficiency ($\phi_{II}$) was assessed over the following 4 days. False-color fluorescence images were acquired every every hour during the 16-hour day period for 4 consecutive days.

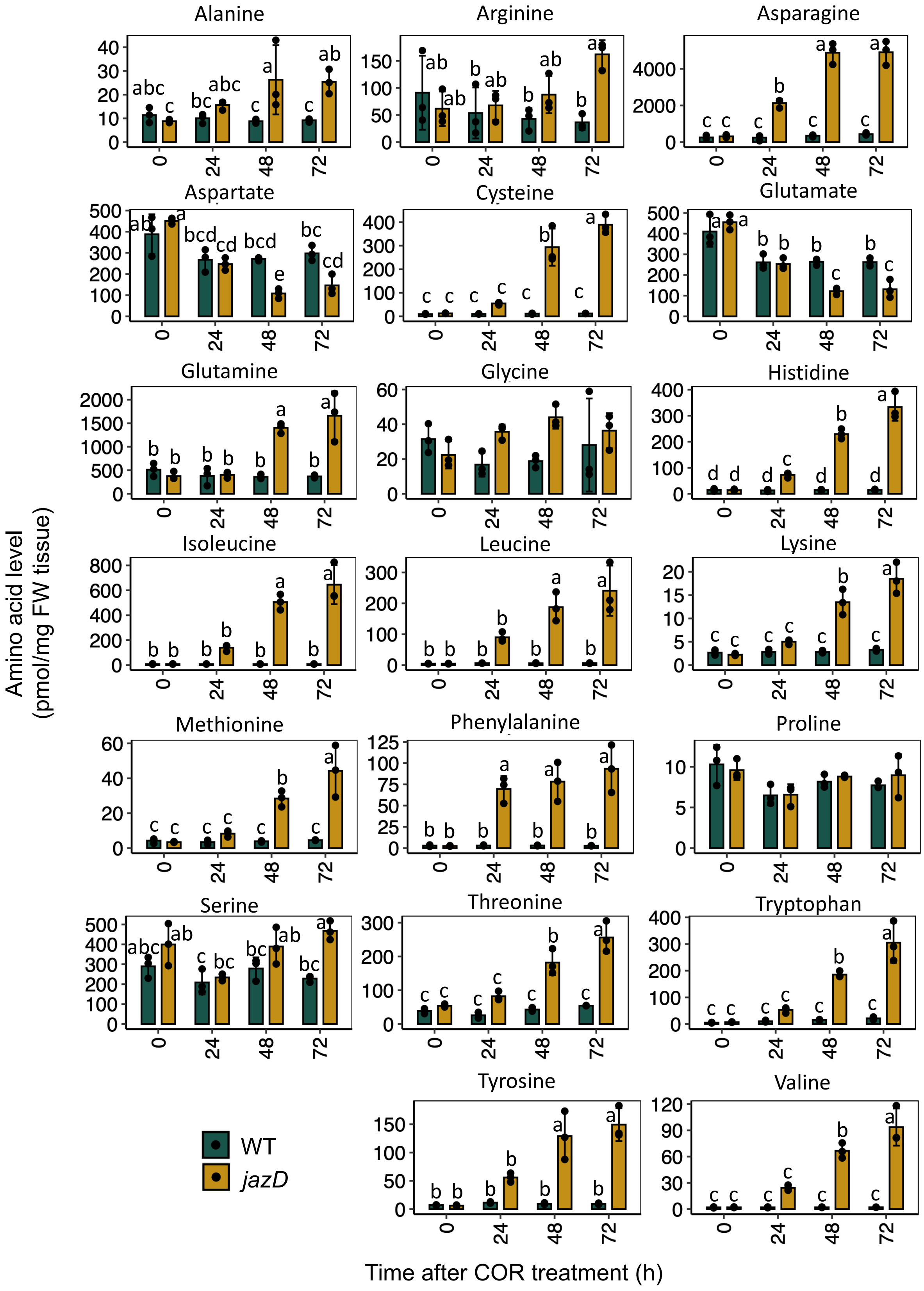


**Supplemental Figure S1.** Coronatine-induced changes in the amino acid content in WT and *jazD* plants. Soil-grown plants were treated with a solution containing 5 µM coronatine. Whole rosettes were harvested at the indicated time points for quantification of amino acids by LC-MS. Different letters represent significant differences at $P<0.05$ with Tukey's honest significant difference (HSD) test.

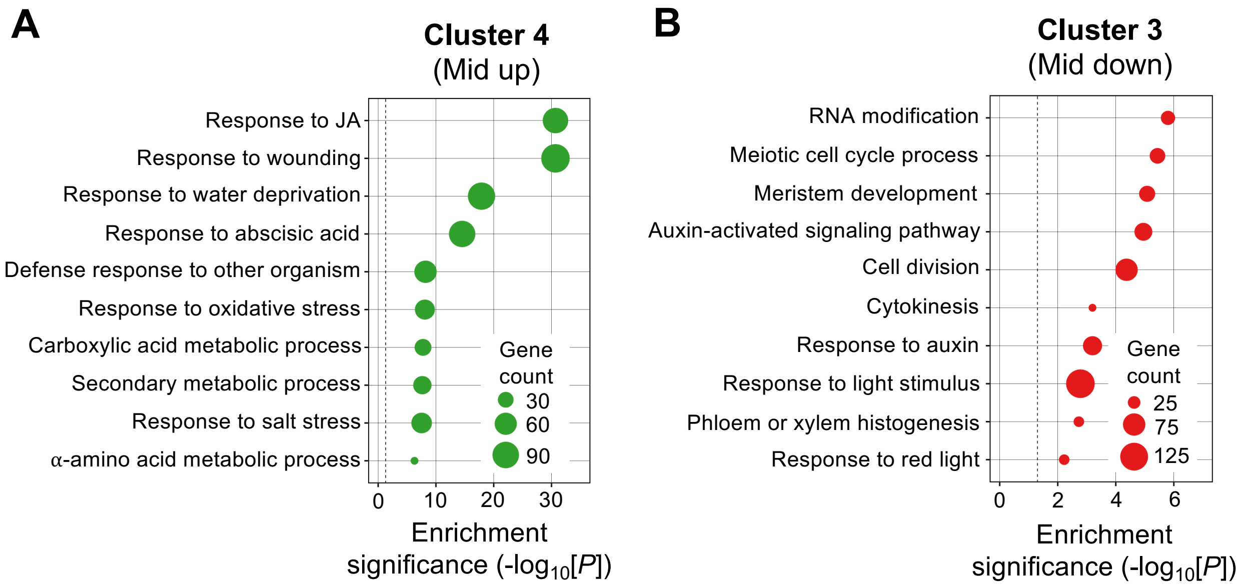


**Supplemental Figure S2.** Global transcriptional response of *jazD* to coronatine treatment. A-B) Gene ontology (GO) analysis (biological processes) was performed with Cluster 4 and 3 genes. Enriched functional categories were determined with DAVID using the hypergeometric test with Benjamini & Hochberg's false discovery rate (FDR) correction.

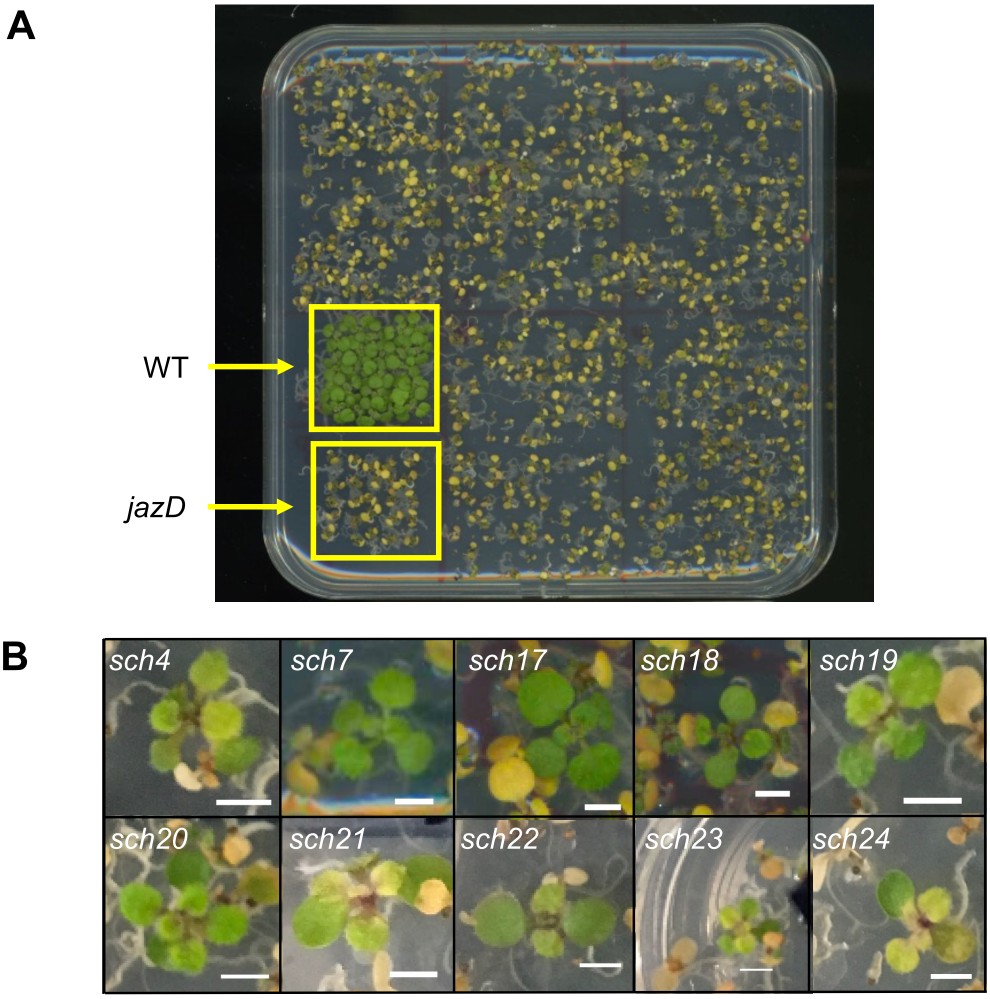


**Supplemental Figure S3.** Identification of coronatine-resistant *sch* mutants. A, Ethyl methanesulfonate (EMS)-mutagenized *jazD* seeds ($M_2$ generation) were germinated on agar medium. Seeds from multiple, independent $M_2$ families were plated in batches of approximately 100 seeds per family. Batches of wild-type Col-0 (WT) and parental *jazD* seeds were plated as controls (lower left, yellow boxes). Eight days after germination, all seedlings were sprayed with a solution containing 1 uM coronatine. Treated seedlings were visually screened for resistance (i.e., reduced chlorosis) four days later, at which time the plate was photographed. B, Photographs of coronatine-resistant *sch* mutants ($M_2$ generation) identified in 10 independent $M_2$ families. Scale bars, 2 mm.

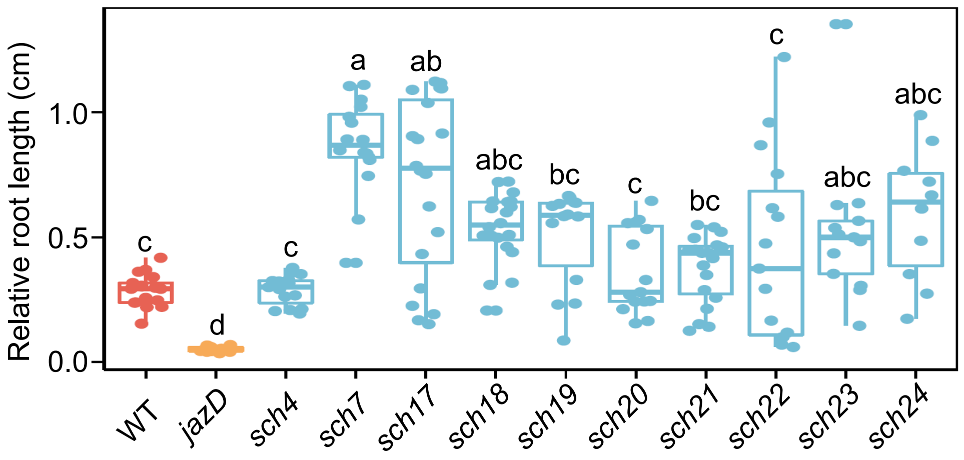


**Supplemental Figure S4.** *sch* suppressor lines have reduced sensitivity to methyl jasmonate (MeJA)-induced root growth inhibition. Seedlings ($M_3$ generation) were grown for 10 days on agar medium supplemented or not (mock) with 10 uM MeJA. Relative root length was determined by normalizing the root length on MeJA-containing medium to that of mock-treated roots. Box plots show the mean and interquartile range, with whiskers extending to the minimum and maximum values but not further than 1.5x the interquartile range (n = 10-20 seedlings per genotype). Letters indicate significant differences at $P<0.05$ according to Tukey's honestly significant difference (HSD) test.

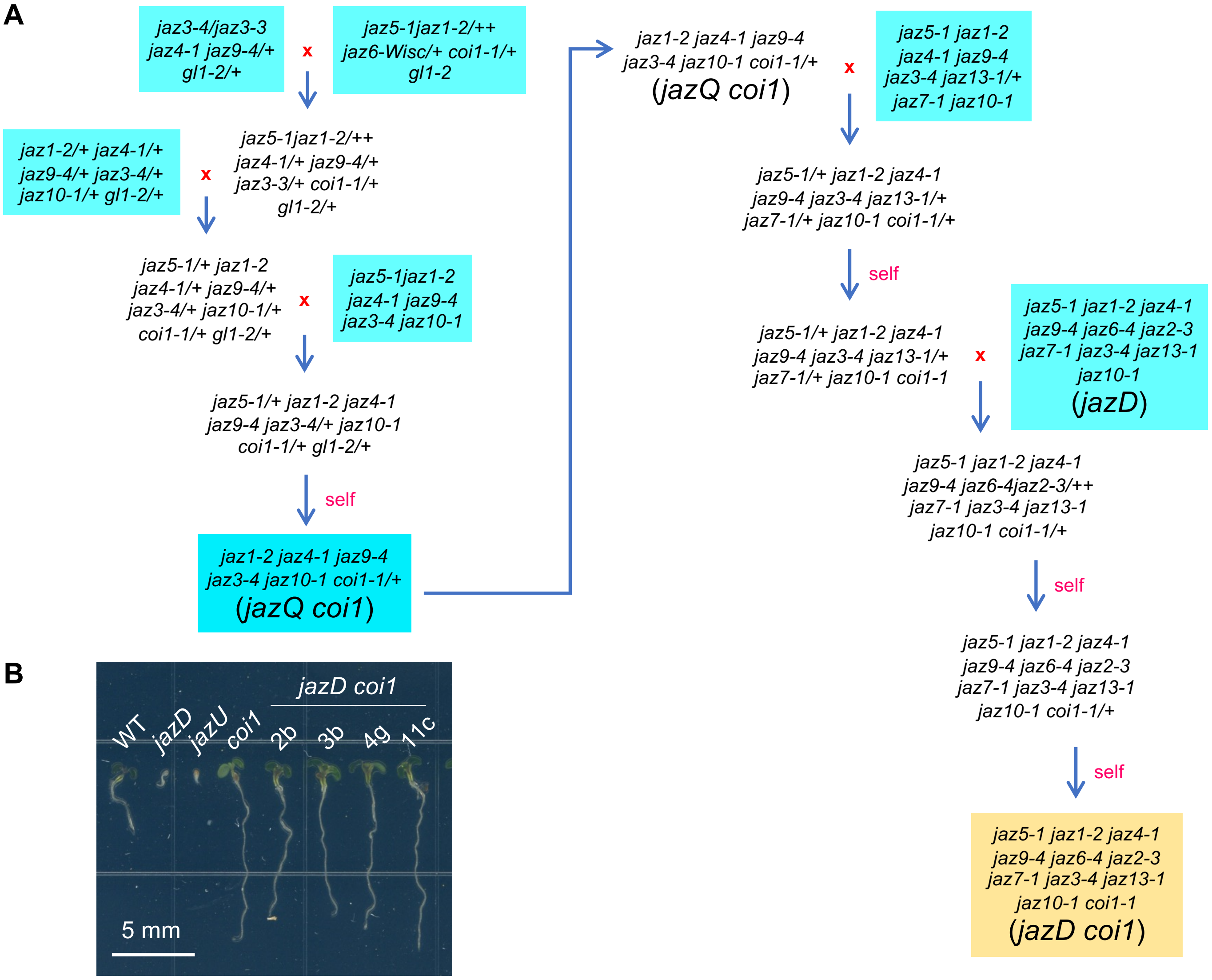


**Supplemental Figure S5**. Construction of *jazD coi1* line for genetic complementation analysis. A, Pedigrees for construction of *jazQ coi1* and *jazD coi1*. Red 'x' and red 'self' indicate cross-pollination and self-pollination, respectively. Mutants in blue shade were reported previously (Major et al., 2017; Guo et al., 2018b). B, Representative seedlings of WT, *jazD*, *jazU*, *coi1*, and independent lines of *jazD coi1* grown on plates supplemented with 10 µM MeJA. *jazD coi1* line 11c was used for crossing to *sch* mutants.

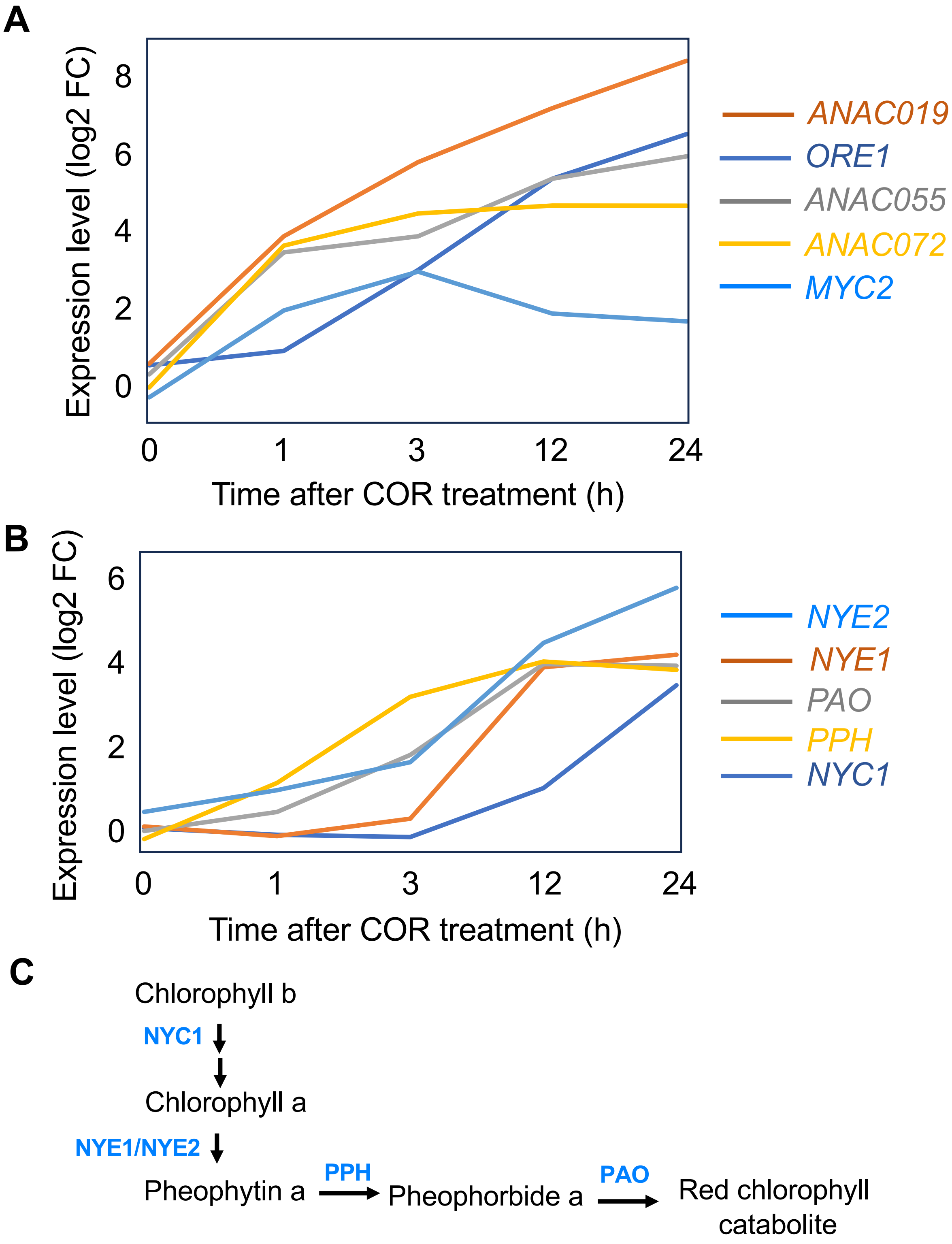


**Supplemental Figure S6**. Coronatine-induced expression of genes encoding select transcription factors (A) and chlorophyll catabolism (B) genes in *jazD*. Gene expression levels are plotted from RNA-seq data (Table S1) and represent the log2 fold-change (FC) in mRNA abundance in the COR-treated sample relative to a time-matched mock control. C, Scheme of chlorophyll breakdown pathway. Enzymes ecoded by genes indicated in panel B are shown in blue text.

Supplemental Table S1. Primers used for Sanger sequencing of *sch* suppressor mutants.

| AGI accession (Gene name)[1] | Primer 1[2] | Primer 2[2] | Amplicon length (bp)[3] |
|---|---|---|---|
| **AT2G39940 (COI1)** | TAAGAACAAGAAATCAAAGT | CTCATATATTGGCTTACTAC | 488 |
| | CATCGACGACACAACATGAA | ATTACAGATCTGCCACTGGA | 642 |
| | GCCTTGTCTCAGATAGAATG | GAGATAAGGGATATGAATGC | 456 |
| | CAATATTGCATTCATATCCC | AAACTTCTACATGACGGAGT | 500 |
| | ACCTTCACAGATACCAGAGA | TTATGCTTGCTATTAAAGCA | 500 |
| | TTGTAGATAAAATCCGGATC | TACTGATGGACTTTTGAGCA | 500 |
| | AACCTCAAAAGCATCGAGCC | ACGGTTGATGATGTCATCGA | 500 |
| | GAACCATCTCCGACACACCA | GTCGTGTAGCTGAGATCTGA | 538 |
| | GCATTCATATCCCTTATCTC | CGGGAAATCCAAATATCTTG | 658 |
| **AT1G32640 (MYC2)** | CTCTTCCTCGATCACCTCGC | CTCTCTGCTTCGACGTGGTT | 1183 |
| | TCCAAATCCATCAACGCCGA | TCCAAATCCATCAACGCCGA | 1189 |
| | AACCACGTCGAAGCAGAGAG | GCGAGGTGATCGAGGAAGAG | 1106 |

[1]AGI accession indicates the locus identifier provided by The Arabidopsis Information Resource (TAIR) database and the Arabidopsis Genome Initiative (AGI). Gene name indicates the common gene symbol.

[2]Primer sequences used for Sanger sequencing are written in the 5' to 3' direction. Multiple primer sets were designed to span the genomic sequence from the 5'UTR to the 3'UTR.

[3]Amplicon length indicates the size of the PCR product.
